# Visualizing the chaperone-mediated folding trajectory of the G protein β5 β-propeller

**DOI:** 10.1101/2023.05.04.539424

**Authors:** Shuxin Wang, Mikaila I. Sass, Yujin Kwon, W. Grant Ludlam, Theresa M. Smith, Ethan J. Carter, Nathan E. Gladden, Margot Riggi, Janet H. Iwasa, Barry M. Willardson, Peter S. Shen

**Affiliations:** Department of Biochemistry, 15 N. Medical Drive East, University of Utah, Salt Lake City, UT, 84112, USA; Department of Chemistry and Biochemistry, C100 BNSN, Brigham Young University, Provo, UT, 84602, USA

## Abstract

The cytosolic Chaperonin Containing Tailless polypeptide 1 (CCT) complex is an essential protein folding machine with a diverse clientele of substrates, including many proteins with β- propeller domains. Here, we determined structures of CCT in complex with its accessory co chaperone, phosducin-like protein 1 (PhLP1), in the process of folding Gβ_5_, a component of Regulator of G protein Signaling (RGS) complexes. Cryo-EM and image processing revealed an ensemble of distinct snapshots that represent the folding trajectory of Gβ_5_ from an unfolded molten globule to a fully folded β-propeller. These structures reveal the mechanism by which CCT directs Gβ_5_ folding through initiating specific intermolecular contacts that facilitate the sequential folding of individual β-sheets until the propeller closes into its native structure. This work directly visualizes chaperone-mediated protein folding and establishes that CCT directs folding by stabilizing intermediates through interactions with surface residues that permit the hydrophobic core to coalesce into its folded state.

## INTRODUCTION

G proteins mediate the transmission of a myriad of extracellular signals across the plasma membrane to the cell interior ^1, 2^. The bridge for signals to traverse the plasma membrane is the G protein-coupled receptor (GPCR), a seven transmembrane helical structure that binds to extracellular ligands and undergoes conformational changes that propagate to the cytoplasmic surface ^3^. Inside the cell, the activated GPCR interacts with the G protein heterotrimer (consisting of Gα, Gβ, and Gγ subunits) to release GDP from the Gα subunit and bind GTP ^4, 5^.

This nucleotide exchange disrupts the association of Gα-GTP with the GPCR and the Gβγ dimer, allowing both Gα-GTP and Gβγ to activate downstream effectors ^6–9^. Gα hydrolyzes GTP to GDP, thereby inactivating the Gα. However, for many Gα isoforms, intrinsic GTP hydrolysis is too slow for physiological responses and requires acceleration by Regulator of G protein Signaling (RGS) proteins ^10, 11^. In neurons, the R7 family of RGS proteins associate with a unique isoform of Gβ (Gβ_5_) to form dimers that interact with Gα and enhance its GTPase activity^12–14^.

To perform their signaling function, G protein subunits must first be folded and then assembled into Gαβ_1-4_γ or Gβ_5_-RGS complexes by molecular chaperones ^15–18^. The primary chaperones for Gβ subunits are the cytosolic chaperonin CCT (also known as T-complex polypeptide 1 ring complex, or TRiC) and its co-chaperone phosducin-like protein 1 (PhLP1) ^15, 17, 19, 20^. CCT is composed of two sets of eight paralogous subunits that form two back-to-back rings, thus creating a barrel-shaped structure with a central folding chamber into which nascent or otherwise unfolded proteins enter and are assisted in folding ^21–23^. Each CCT subunit is an ATPase that binds and hydrolyzes ATP to drive a conformational change that seals off the ends of the barrel and encapsulates the unfolded protein, allowing it to fold in the sequestered environment of the folding chamber ^24–26^. After phosphate release, the ring reopens and enables the folded protein to be released.

CCT has been estimated to fold up to 10% of the cytosolic proteome ^27^, particularly proteins with multiple domains and complex folding patterns ^28^. Gβ subunits belong to the β-propeller family of protein folds, a major class of CCT substrates with important physiological functions^29^. This family of β-propellers contains seven WD40 repeat sequences that fold into structures consisting of seven β-sheets arranged into a radial, propeller-like structure (Figure 1A) ^30–32^. For β-propellers to fold completely, the N- and C-terminal β-strands of the β-propeller must be brought together to form the last β-sheet ^20^. Yet, the mechanism underlying CCT-mediated folding remains unresolved.

**Figure 1.**
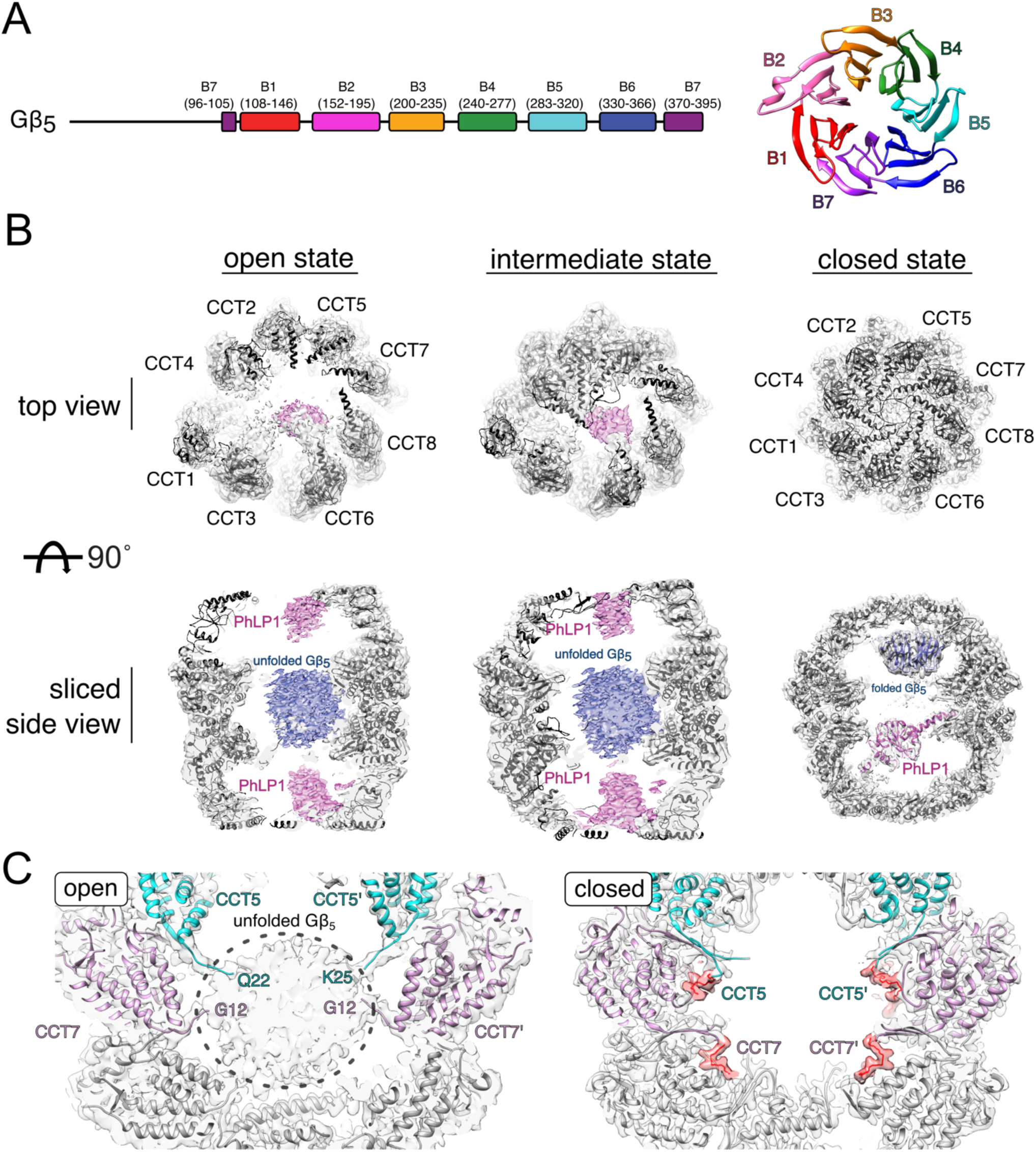
Reconstructions of Gβ_5_-PhLP1-CCT complexes. (A) Domain organization and structure of folded Gβ_5_. B1-7, Blades 1-7. (B) Views of CCT-PhLP1-Gβ_5_ reconstructions in open, intermediate, and closed states computed from a single dataset. CCT, gray; PhLP1, pink; Gβ_5_, blue. (C) Conformational change in hydrophobic N-terminal arms of CCT5 and CCT7 between open and closed states. Density and model of additional ordered N-terminal residues observed in the closed state of CCT5 and CCT7 are colored red.

The CCT co-chaperone, phosducin-like protein-1 (PhLP1), plays an important role in Gβ subunit folding and assembly into Gβγ and Gβ_5_-RGS dimers. PhLP1 binds CCT and helps to fold Gβ_1-4_ and Gβ_5_ and then release them to interact with their Gγ or RGS binding partner ^15, 16, 19, 20^. The vital role of PhLP1 in both Gβγ and Gβ_5_-RGS dimer formation has been confirmed *in vivo*, where deletion of PhLP1 in mouse photoreceptor rods ^33^, cones ^34^, and striatal neurons ^35^ resulted in reduced levels of Gβγ and Gβ_5_-RGS complexes and corresponding disruption of G protein signaling.

To understand the molecular mechanism underlying the roles of CCT and PhLP1 in Gβ_5_ folding, we isolated Gβ_5_ in complex with CCT and PhLP1 in the process of folding and used cryo-EM to determine an ensemble of structures that capture its complete folding trajectory. The structures reveal how CCT and PhLP1 orchestrate the Gβ_5_ folding trajectory from an unfolded molten globule state to its fully folded β-propeller. Partially folded intermediate states are stabilized by hydrophilic interactions between surface residues of Gβ_5_ and the apical domains of CCT that allow the hydrophobic core of Gβ_5_ to coalesce into its β-propeller structure.

## RESULTS

### Reconstructions of CCT-PhLP1-Gβ_5_ complex

CCT-PhLP1-Gβ_5_ particles were isolated directly from HEK293T cells virally transduced with His6-tagged human PhLP1 and Strep-tagged human Gβ_5_ (Figure S1A). A two-step affinity purification scheme was used to first isolate PhLP1 complexes on a Ni^2+^-chelate column, followed by Gβ_5_ complexes on a StrepTactin column. Coomassie blue stained SDS-PAGE showed strong bands corresponding to the expected molecular weights of CCT subunits, along with a band resulting from the co-migration of PhLP1 and Gβ_5_. Immunoblot analysis confirmed the presence of PhLP1, Gβ_5_ and CCT in the purified sample. ATPase assays showed the complex hydrolyzed ATP at a rate comparable to CCT alone, indicating that the complex was functional (Figure S1B). Previous studies demonstrated that substrate folding is initiated by inducing a conformational change from an open to a closed state of CCT in an ATP-dependent manner ^36, 37^. In order to capture CCT complexes in the act of folding Gβ_5_, we incubated CCT-PhLP1-Gβ_5_ particles with ATP and aluminum fluoride to generate the transition state analogue ADP-AlF_x_ and induce the closed form. The inclusion of ATP in the initial reaction permits at least one round of ATP hydrolysis, while the hydrolyzed nucleotide is presumably stabilized by AlFx. Native gel analysis after nucleotide treatment confirmed a conformational shift in CCT, consistent with a transition from the open to the closed form as the complex becomes more compact ^26, 38^ (Figure S1C).

Purified CCT-PhLP1-Gβ_5_ particles were used for single particle cryo-EM imaging. Image classification revealed the enrichment of three distinct conformational states of CCT, including open, intermediate, and closed states with resolutions at 3.4 Å, 3.7 Å and 2.4 Å, respectively (Figures 1B and S2). The intermediate state for CCT shows CCT subunits with higher ATPase activity (CCT2, 4, 5) adopting a closed conformation, while those with lower ATPase activity remain in the open conformation, leaving the CCT barrel partially closed. Both the open and intermediate states contained additional densities within CCT at the apical and equatorial regions at poor local resolution. The size of the equatorial density roughly matched the expected volume of Gβ_5_, and its position between the CCT rings was consistent with other unfolded substrates that have been reported previously ^36, 37, 39^. Our reconstruction also reveals N-terminal extensions contributed by both copies of CCT5 and CCT7 that tether the unfolded substrate within the center of the barrel (Figure 1C). These N-terminal regions are enriched in hydrophobic residues, suggesting that hydrophobic contacts partially contribute to the stabilization of the unfolded Gβ_5_ substrate. These observations indicate that the open and intermediate states of CCT hold Gβ_5_ in its cavity near the equatorial domain through interactions mediated by CCT5 and CCT7, but the substrate is held in a compact molten globule state.

**Figure 2.**
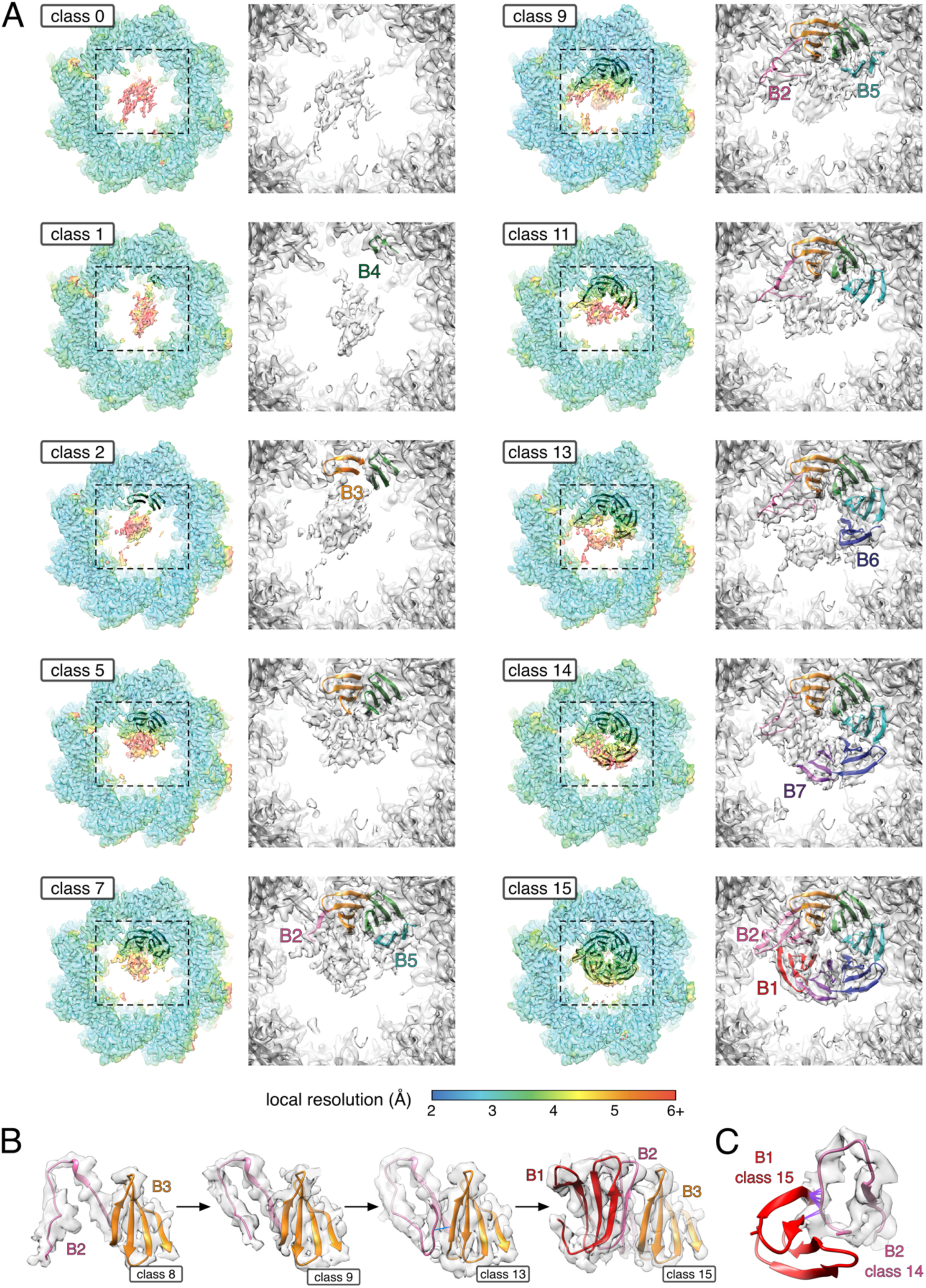
The folding trajectory of Gβ_5_. (A) Focused views of eight folding intermediates reveal the stepwise folding process of Gβ_5_ from a molten globule state (class 0) to a complete β-propeller (class 15). Well resolved Gβ_5_ regions were determined by local resolution heat maps (left) and models are shown as colored ribbons (right), beginning in class 1. Unmodeled densities represent unfolded/disordered segments of Gβ_5_ (red densities in local resolution maps). See Figure S6 and Supplemental Movie 1 for views of all sixteen classes. (B) Focused views of four classes demonstrate that B2 is adopts a range of extended conformations during Gβ_5_ folding. This extended conformation is stabilized in part by hydrogen bond interactions with B3 (class 13). In the final step (class 15), B2 refolds into a canonical β-sheet and creates space for B1 to form and complete the β-propeller. (C) Clashes (purple lines) between B2 in class 14 and B1 in class 15 indicate that B2 sterically hinders B1 folding.

The size and shape of the two densities associated with the apical region of CCT6 in each ring are consistent with the compactly folded thioredoxin domain (TXDN) of PhLP1, while no density matched with the PhLP1 N-terminal domain, which is known to form extended α-helices and random coils ^37, 40, 41^. A difference map between this reconstruction and one lacking PhLP1 indicate that the extra densities belong to PhLP1 (Figure S1D and S3).

To verify the positions of Gβ_5_ and PhLP1 in the CCT open state, we performed crosslinking mass spectrometry (XL-MS) of purified CCT-PhLP1-Gβ_5_. Crosslinking distances were minimized when Gβ_5_ was positioned in the mass between the CCT rings (Figure S4). For PhLP1, no crosslinks were observed between the PhLP1 thioredoxin domain and CCT, but several were identified between the N-terminal domain and CCT4 and 5 in the equatorial and intermediate regions, indicating that this domain extends deeper into the CCT folding chamber and is not the PhLP1 density associated with the apical domain of CCT6. Based on these results, it appears that unfolded Gβ_5_ is tethered to the center of CCT in its open form by extended N-terminal arms from CCT5 and CCT7, and the complex is capped at the apical ends by the PhLP1 thioredoxin domain with the PhLP1 N-terminal domain extending into the CCT cavity toward Gβ_5_.

### Gβ_5_ and PhLP1 become ordered in the closed state of CCT

In contrast to the open and intermediate states, the densities for Gβ_5_ and PhLP1 became well-resolved in the closed state of CCT (Figures 1B and S2). Gβ_5_ and PhLP1 occupy opposing ends of the closed CCT chamber in different positions compared to the open state (Figure 1B). Most notably, Gβ_5_ shifts positions from being tethered between the CCT rings in the open state and moves to the apical region of CCT2, 5, and 7 in the closed state. Interestingly, the N-terminal arms of CCT5 and CCT7 fold back onto the CCT inner wall in the closed CCT structure (Figure 1C). These conformational differences between the open and closed states indicate a mechanism in which the N-terminal arms hold unfolded substrate in the center of the open CCT barrel, followed by the withdrawal of the arms upon CCT closing to release substrate into the folding chamber.

### The folding trajectory of Gβ_5_

In the closed state, the consensus map of Gβ_5_ contains density of a partially folded β- propeller with variable local resolution (Figure S2F). The presence of partially ordered Gβ_5_ density prompted us to investigate if the reconstruction represented a mixture of different conformational states. In order to probe for conformational heterogeneity, we applied 3D variability analysis as implemented in cryoSPARC ^42^ using a focused mask over the folding chamber of closed CCT (Figure S5). The principal component exhibiting the greatest degree of variability revealed heterogeneity within Gβ_5_ densities. Subsequent focused classification over Gβ_5_ revealed an ensemble of 16 distinct classes that represent a range of Gβ_5_ folding intermediates at variable local resolutions (Figures 2A and S6). The classes were assembled into a progression of Gβ_5_ folding based increasing completeness and improved local resolution of Gβ_5_ density (classes 0-15), beginning with a molten globule state with poor local resolution (class 0) to a reconstruction with high local resolution that enabled the modeling of a fully folded β-propeller (class 15) (Supplemental Movies 1, 2). Each class represents a distinct snapshot of Gβ_5_ folding, and models of folding intermediates were built into charge density that displayed sufficient local resolution for the separation of β-strands. The ordering of classes reveals a trajectory that begins with the migration of unfolded Gβ_5_ from the equatorial to apical end of CCT, and the ordering of snapshots reveals the sequence of folding for each of the seven blades of the β-propeller fold (B1-B7). Throughout the trajectory, charge density at poor local resolution is observed in multiple positions, as if the densities were shifting as a molten globule to sample various conformations without adopting an ordered state. Models were not built into these unstructured portions, although classes 1-14 show that they are connected to well-resolved segments of Gβ_5_.

In the first snapshot, Gβ_5_ is poorly resolved as a molten globule in a position as if it were just released from the N-terminal arms of CCT5 and CCT7 upon CCT closure (class 0). In the ensuing class, the globule migrates towards the apical region and the outermost β-strand of B4 initiates contact with CCT5 (class 1). This blade can be attributed to B4 because it maintains this position in all the classes up to the completely folded β-propeller (class 15) in which the position of each blade can be unambiguously modeled based on the fit of the sequence into the charge density map, including the 3_10_ helix that is unique to B2. The class 1 snapshot is followed by four classes that reveal the concurrent formation of B3 and B4 into complete four-stranded β-sheets (classes 2-5). Following B3 and B4, folding progresses radially around the propeller such that B2 and B5 begin to form (classes 6-11), followed by B6 (classes 12, 13), and B7 (class 14). Among classes 6 through 14, B2 does not assemble into a β-sheet and is instead stabilized in a range of extended conformations while B5, B6, and B7 fold sequentially. In the final class, B2 is rearranged into its native conformation and the formation of B1 seals the edges of the β-propeller and completes the folding process (class 15).

### Folding is guided by stabilizing intra-molecular interactions within Gβ_5_

As described above, B2 is observed in a range of extended conformations during the sequential folding of B5, B6, and B7 (Figure 2B). These classes account for approximately 45% of particles in the CCT closed state, which suggests that this incomplete B2 intermediate is stabilized during Gβ_5_ folding. In this conformation, B2 is partially stabilized through the hydrogen-bond contacts with the β-sheet of the neighboring B3, and this position sterically obstructs the formation of B1 (Figure 2C). Following the formation of B7, B2 undergoes a conformational change and reorganizes into its native β-sheet, which enables B1 to fold and complete the folding process of the β-propeller. The stabilization of B2 during the folding trajectory of Gβ_5_ suggests a mechanism that delays the temporal folding of B1. By occluding the space needed for B1 formation, this ensures that all other blades have folded before B1 seals the edges to complete the β-propeller.

### Folding is initiated and guided by specific interactions between CCT and Gβ_5_

The first interaction between Gβ_5_ and CCT is mediated by the formation of the outermost β-hairpin that initiates the folding of B4 (class 1, Gβ_5_ residues 267-274). Interestingly, R269 within this hairpin is conserved in all Gβ proteins and is the only surface-exposed basic residue within a ∼6,450 Å^2^ patch of folded Gβ_5_ (Figure 3A). This residue forms a complementary electrostatic interface with a strong acidic patch in CCT5 (Figure 3B). The interaction appears highly specific as the charge density map in class 1 reveals an electrostatic interaction between R269 of Gβ_5_ (B4) and E256 of CCT5 (Figure 3C). The β-hairpin also appears to be stabilized by a hydrogen-bond between D267 of Gβ_5_ with the indole nitrogen of W324 (CCT5) and a hydrophobic contact between V274 of Gβ_5_ with the same tryptophan residue (Figure 3C). No other interactions between Gβ_5_ and CCT were observed in this class, suggesting that the initiation of folding is driven by the three contacts between Gβ^5^ and CCT5 (R269-E256, D267-W324, and V274-W324).

**Figure 3.**
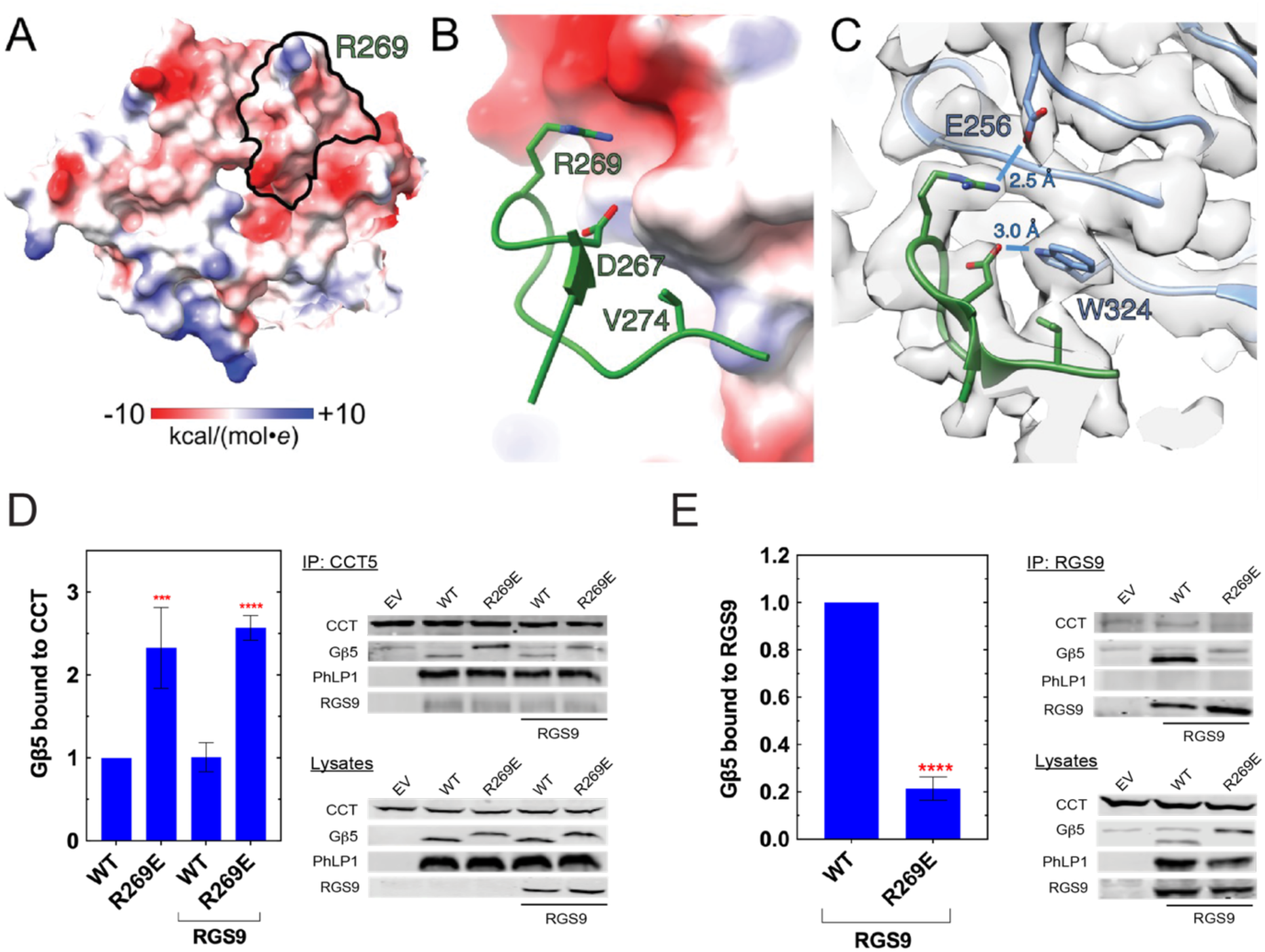
Gβ_5_ folding is initiated by specific contacts between B4 and CCT5. (A) The electrostatic potential map of Gβ_5_ reveals a single surface-exposed basic residue (R269) within B4 (outlined) and surrounding regions. Red, surface-exposed acidic residues; blue, surface exposed basic residues. (B) The electrostatic potential map of CCT5 reveals a surface-exposed negatively charged pocket that accommodates R269 of Gβ_5_. The interaction is also further stabilized by additional contacts involving D267 and V274. (C) A series of electrostatic, hydrogen-bond and hydrophobic interactions stabilize the position of the outermost β-hairpin of B4 with CCT5. (D) The R269E mutation increases Gβ_5_ association with CCT with or without RGS9. Note that the R269E charge reversal slows the migration of the variant in SDS gels compared to Gβ_5_ WT causing R269E to co-migrate with a non-specific band in the Gβ_5_ blots. To correct for this, the intensity of the non-specific band in the empty vector lane was subtracted from the R269E band in the quantifications. (E) The R269E mutation reduces Gβ_5_ association with RGS9. Bars represent the average ± standard deviation normalized to the WT Gβ_5_ sample (n = 5). Red *** p < 0.001, **** p < 0.0001 in t-tests compared to WT Gβ_5_.

To further assess the contribution of the salt bridge between Gβ^5^ R269 and CCT5 E256 to Gβ_5_ folding, we expressed a Gβ^5^ R269E variant in cells and measured its ability to bind CCT with or without RGS9 (Fig. 3D). Gβ_10_ R269E showed more than a 2-fold increase in binding to CCT compared to WT in a manner that was independent of RGS9. This increase in CCT binding is consistent with a model in which substrate folding is impaired, leading to the failure of substrate to release from CCT. In further support of this model, Gβ_5_ R269E binding to RGS9 was decreased 5-fold compared to WT (Fig. 3E). These results indicate that the R269E substitution disrupts CCT-mediated Gβ^5^ folding, resulting in failure to mature into a state that can release from CCT to interact with its cognate RGS9 binding partner. These findings therefore support the idea that the salt bridge between Gβ^5^ R269 and CCT5 E256 initiates folding.

Additional electrostatic and hydrogen-bonding interactions that stabilize folding intermediates are observed in subsequent classes. Specifically, B4 is further anchored to CCT5 through hydrogen-bonds between S248 of Gβ_5_ and K259 of CCT5 (Figure 4A). As other blades fold radially around B4, additional interactions are observed between B5 (R311) and CCT7 (E245/D297) (Figure 4B). Hydrogen-bonding interactions are also observed between main chain carbonyls in B3 (L229, L330) and the side chain of H314 in CCT2 (Figure 4C). Additional hydrogen bond interactions are observed between B5 with CCT5 and between B3 with CCT4 in class 15 (Figure 4D). In contrast, no Gβ_5_-CCT interactions are observed between B1, B2, B6, and B7. Taken together, the stabilization of B3 with CCT2, B4 with CCT5, and B5 with CCT7 through polar interactions likely seeds the folding of Gβ_5_ to enable the sequential formation of each blade, beginning with B4 (Figure 4D). This extensive hydrogen bond and electrostatic interaction network in the Gβ_5_-CCT interface indicates that CCT stabilizes Gβ_5_ folding through hydrophilic interactions with surface residues, leaving the hydrophobic core free to collapse into its folded conformation.

**Figure 4.**
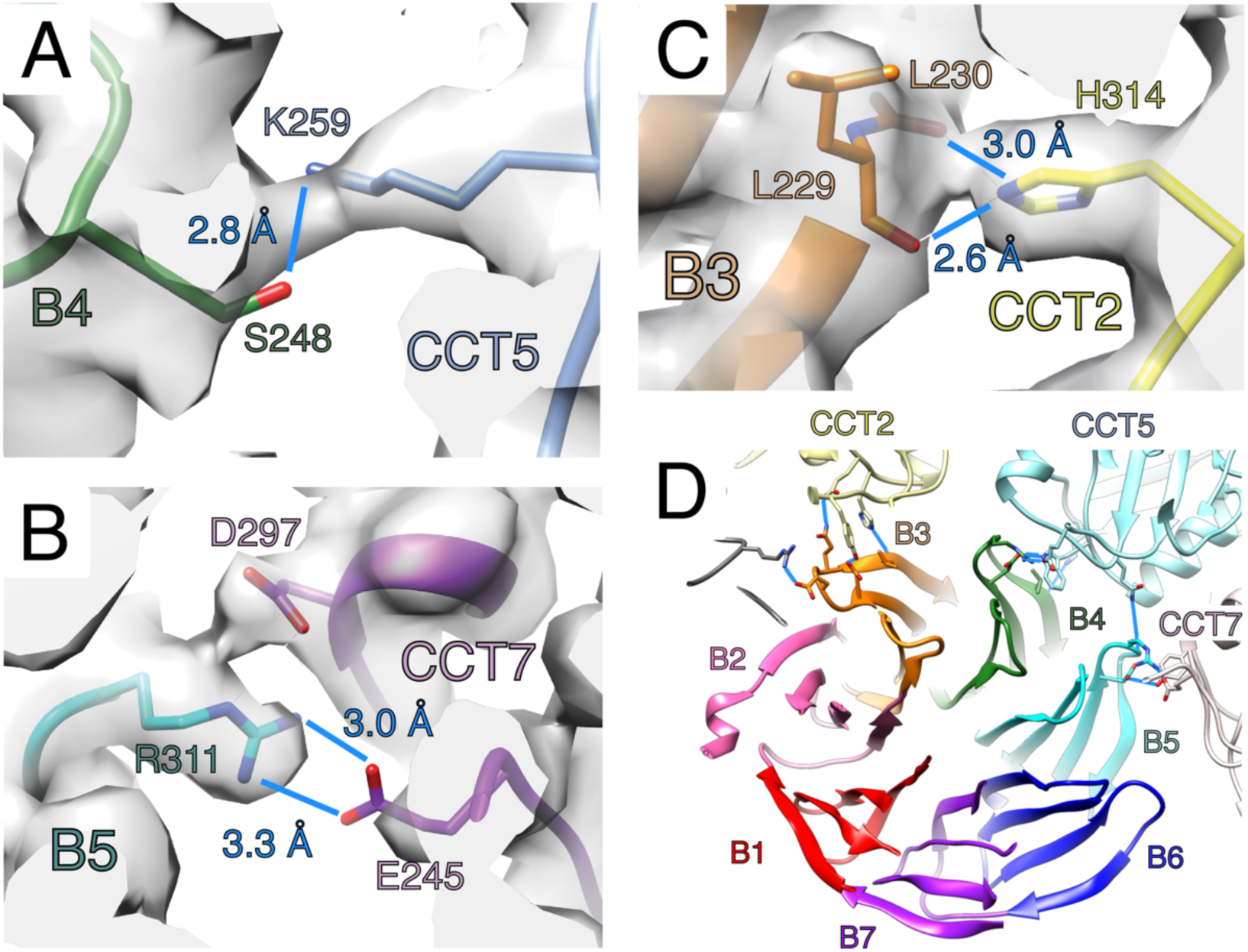
Gβ_5_ folding is stabilized by a network of interactions between B3-5 and CCT2, 5, and 7. Focused views of specific CCT interactions with Gβ_5_ folding intermediates. (A) Hydrogen bond interaction observed between S248 of Gβ_5_ and K259 of CCT5 in class 5. (B) Hydrogen bond formed between CCT2 (H314) and backbone carbonyls of B3 (L229, L230) in class 2. (C) Salt bridge formed between B5 (R311) and CCT7 (E245) in class 6. All interactions are maintained in subsequent classes. (D) Hydrogen-bond network formed between B3, B4, and B5 with specific CCT subunits in the fully folded state of Gβ_5_ (class 15). All hydrogen bond interactions are shown as blue lines.

### PhLP1 spatially confines Gβ_5_ to a defined region within CCT

The two copies of the PhLP1 co-chaperone also relocate when CCT closes. One copy is no longer found in the structure, presumably because it has dissociated from CCT, while the other copy dissociates from its position on the apical domain of CCT6 to occupy the folding chamber opposite of Gβ_5_ (referred to as the CCT′ chamber). The local resolution permitted building and refining an atomic model of PhLP1 encompassing residues 55-93 and 136-301, which include two extended helices (H1 and H2), followed by a conserved thioredoxin domain (TXDN), and ending with a structured C-terminal tail (Figures 5A and S7A). Contacts between PhLP1 and CCT are mediated by a network of polar interactions that span a large surface area across multiple CCT subunits (Figure 5B). H1 (residues 57-68) lines against CCT1′, 3′, and 4′, while the C-terminal tail of PhLP1 is highly acidic and extends through a deep, positively charged groove between CCT1′ and CCT4′ and into the exterior of CCT. H2 and the TXDN are well resolved, and the linker between these two regions associates with CCT1′, CCT3′, and CCT6′ through H-bonds.

**Figure 5.**
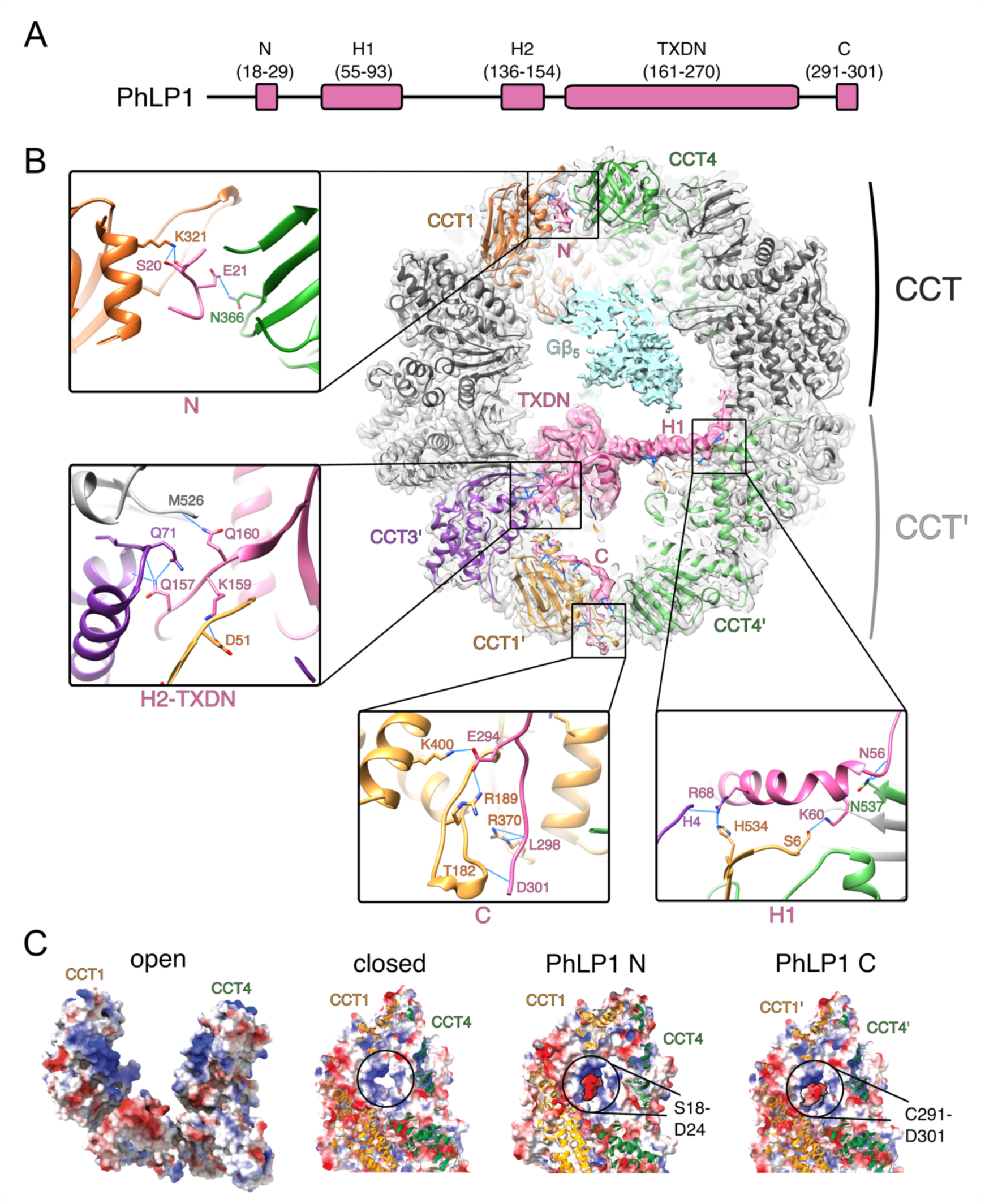
PhLP1 forms an extensive network of interactions with CCT. (A) Domain organization of PhLP1. N, negatively charged N-terminal motif; H1, helix 1; H2, helix 2; TXDN, thioredoxin domain; C, negatively charged C-terminal motif. (B) Cut-away view of PhLP1 (pink) interactions with CCT. Select zoomed-in views of PhLP1-CCT interactions shown: N-terminal negatively charged residues with CCT1 and CCT4 (top left), H2-TXDN linker with CCT3′, CCT1′, and CCT6′ (middle left), C-terminal negative charged residues with CCT1′ (bottom middle), and H1 with CCT1′ and CCT4′ (top right). Hydrogen bond interactions between labeled residues shown as blue lines (distances measured between 2.7-3.3 Å). Unfolded Gβ_5_ density is colored light blue. (C) The closed state of CCT at the CCT1/4 interface is stabilized by complementary charge interactions from the negatively charged N- and C-motifs in PhLP1 (electrostatic surfaces shown, PhLP1 surfaces are outlined).

PhLP1 shares sequence and structure homology to the PhLP2A co-chaperone, the latter of which is best characterized for its role in CCT-mediated folding of actin. Interestingly, PhLP1 and PhLP2A interact with different components of CCT and are positioned in different regions relative to substrate (Figure S7B). While PhLP2A interacts with CCT primarily via its TXDN domain ^37^, the same domain in PhLP1 does not appear to directly contact CCT. In contrast, the position of PhLP1 is stabilized through an extensive network of interactions that span multiple CCT subunits, thereby increasing the avidity of PhLP1 binding and restraining its position to a defined region within CCT. The interactions between co-chaperones with substrate are also dissimilar. PhLP2A contributes an N-terminal α-helix that extends into the other folding chamber to bind actin ^37^, whereas an ordered interaction between PhLP1 and Gβ_5_ is not observed.

Density for PhLP1 becomes disordered towards the N-terminus of H1 as it traverses into the substrate-containing half of CCT, in proximity to unfolded substrate (Figure 5B). However, density emerges at the CCT1/4 interface approximately 11 Å from the unfolded Gβ_5_ density. The interface is structurally equivalent to the opposing CCT1′/4′ interface that accommodates the acidic PhLP1 C-terminal tail. Interestingly, a series of acidic and phosphorylated residues near the N-terminus of PhLP1 (residues 18-29) ^16^ are predicted to form an α-helix, and structural prediction algorithms indicate that this helix interacts with CCT1 in a position that overlaps with the observed density (Figure S7C). Based on this modeling, it appears that PhLP1 N- and C-termini span between the positively charged CCT1/4 interface in one folding chamber to the equivalent groove in the opposite chamber (Figure 5B). This conformation would indicate that PhLP1 passes around unfolded Gβ_5_, although no direct interaction between PhLP1 and Gβ_5_ is observed likely due to poor local resolution of the unstructured substrate. The PhLP1-CCT interactions also appear to stabilize the CCT1/4 interface, which otherwise would be expected to encounter charge repulsion from the strongly basic patches in CCT1 and 4 (Figure 5C).

Interestingly, this same interface anchors the acidic E-hook of tubulin, which may explain why tubulin does not need a PhLP co-chaperone within the CCT chamber to facilitate folding ^36^.

To understand the functional significance of this extended PhLP1 structure, we prepared a series of PhLP1 truncations and measured their effect on Gβ_5_ binding to CCT and the formation of Gβ_5_-RGS9 dimers in co-immunoprecipitation experiments (Figure 6). Full-length PhLP1 increased Gβ_5_ binding to CCT several fold compared to Gβ_5_ alone, as expected ^19^. In contrast, co-expression of RGS9 with PhLP1 led to a reduction in binding to CCT as Gβ_5_ releases upon dimer formation with RGS9 (Figures 6B). The progressive truncation of PhLP1 N-terminal residues up to PhLP1 Δ129 caused a striking increase in Gβ_5_ binding above that of full length PhLP1 and a proportionate decrease in Gβ_5_ binding to RGS9 (Figures 6B, C). The effects of the PhLP1 Δ136 truncation were reduced presumably because this truncation disrupted binding to CCT (Figure 6B, PhLP1 blot), which is consistent with previous work showing that PhLP1 residues 130-136 contributed to binding the open form of CCT ^43^. In contrast to the N-terminal deletions, the truncation of the C-terminal tail of PhLP1 that extends from the thioredoxin domain (residues 277-301) did not change Gβ_5_ binding to CCT or RGS9 compared to full-length PhLP1, suggesting that the interaction of the PhLP1 C-terminal tail at the CCT1′/4′ interface does not influence Gβ_5_ folding by CCT in the opposite chamber. Collectively, the increased Gβ_5_ binding to CCT and the increased Gβ_5_-RGS9 dimer formation with full-length PhLP1, coupled with the increases in Gβ_5_ binding to CCT and decreases in Gβ_5_-RGS9 dimer formation with the N-terminal truncations, suggest a dual effect of PhLP1 on Gβ_5_ folding in which the thioredoxin domain enhances Gβ_5_ binding to CCT while the N-terminal extension mediates its release to interact with RGS9 (see Discussion).

**Figure 6.**
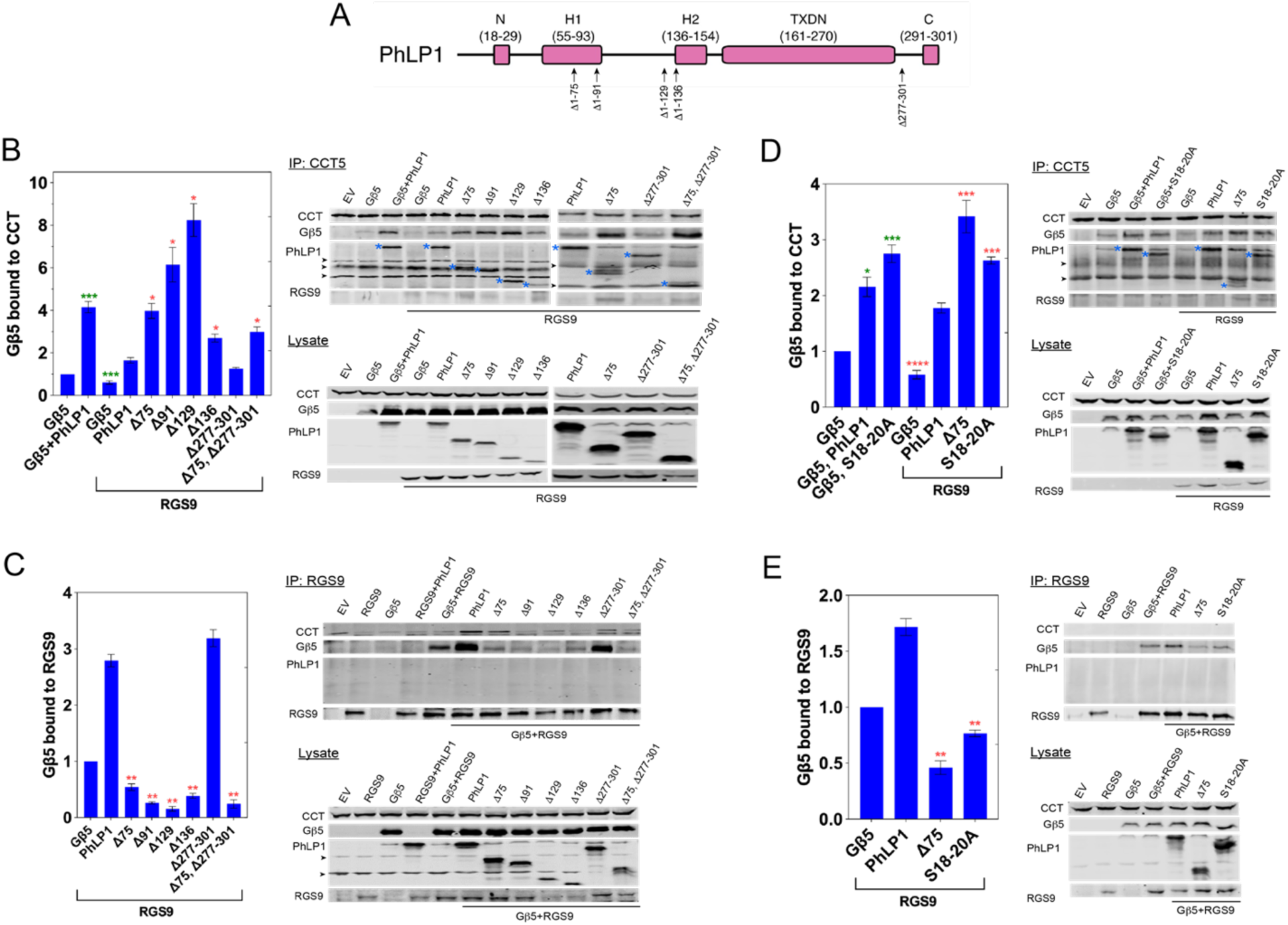
Role of PhLP1 domains in Gβ_5_ binding to CCT and RGS9. (A) Domain organization of PhLP1 with truncation sites indicated. (B) Effects of co-transfection of full length PhLP1 and several truncation variants on Gβ_5_ co-immunoprecipitation (co-IP) with CCT or (C) RGS9. (D) Effects of co-transfection of the CK2 phosphorylation site variant S18-20A on Gβ_5_ co-IP with CCT or (E) RGS9. Bars represent the average ± standard deviation normalized to the Gβ_5_ alone sample (n = 3). Blue asterisks * mark co-IP bands of WT PhLP1 or PhLP1 truncations in CCT IPs with arrowheads indicating non-specific bands. Green asterisks * p < 0.05, *** p < 0.001 in t-tests compared to Gβ_5_ alone. Red asterisks * p < 0.05, ** p < 0.01, *** p < 0.001, **** p < 0.0001 in t-tests compared to the Gβ_5_, PhLP1, RGS9 sample.

The N-terminal extension of PhLP1 harbors a protein kinase CK2 phosphorylation site at three consecutive serine residues (S18-20) that is essential for Gβγ assembly ^15, 16^. Interestingly, these residues are part of the acidic stretch that interacts with the basic interface between CCT1 and 4. To assess the contribution of phosphorylated S18-20 to Gβ_5_ folding, we measured the effect of a PhLP1 S18-20A alanine substitution variant on Gβ_5_ binding to CCT and RGS9 (Figures 6D-E). This variant increased Gβ_5_ binding to CCT and decreased binding to RGS9 almost to the same extent as the N-terminal Δ75 truncation, indicating that the phosphorylated S18-20 contributes significantly to Gβ_5_ folding and Gβ_5_-RGS dimer formation.

## Discussion

The CCT complex is an essential and abundant molecular machine that is responsible for folding a diverse clientele of substrates, yet the underlying mechanism by which it facilitates folding has remained unresolved. Here, we isolated CCT-PhLP1 particles in the act of folding Gβ_5_, a core component of the R7 family of RGS signaling complexes, and performed cryo-EM and image processing to reveal its folding mechanism. The results enabled detailed molecular modeling for Gβ_5_ folding from its unfolded, molten globule state to a fully folded β-propeller (Figures 7A, B and Supplemental Movies 1, 2). Our models establish that unfolded Gβ_5_ initially associates with the open form of CCT in a space between the rings and is tethered by interactions with the hydrophobic N-terminal tails of CCT5 and CCT7 from both rings. ATP binding and progression to the ATP hydrolysis transition state drives a conformational change that brings the apical domains together to close the folding chamber and withdraws the N-terminal tails against the equatorial domains, thus releasing the unfolded Gβ_5_ from between the CCT rings into the folding chamber.

**Figure 7.**
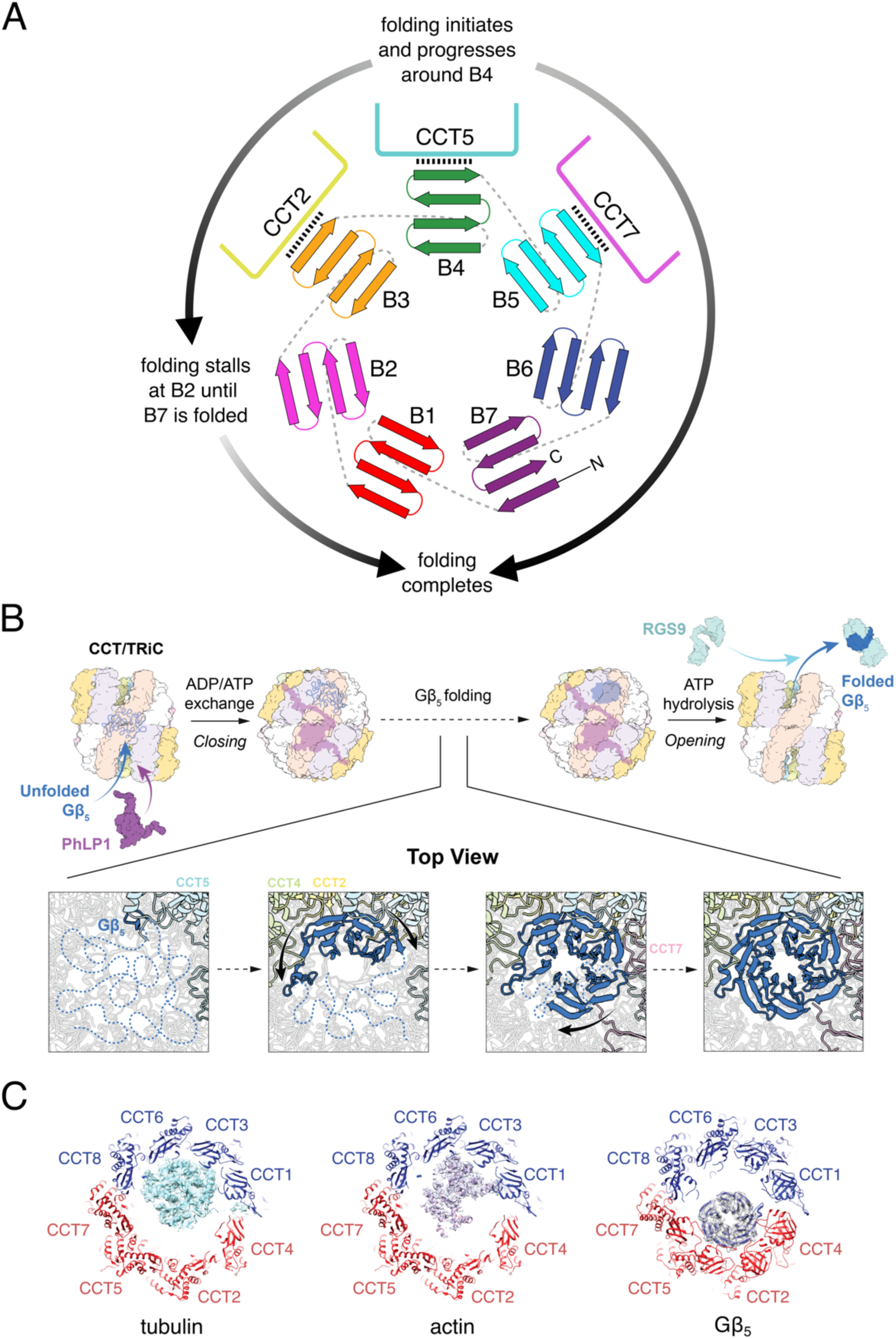
Model of Gβ_5_ folding by CCT. (A) Beta-sheet topology of Gβ_5_ depicting directionality of folding and interactions between B3, B4, and B5 with CCT2, 5, and 7, respectively. Schematic is not drawn to scale. (B) Model figure depicting key steps in CCT mediated folding, from encapsulation of the unfolded protein in the open state, to progressive folding in the closed state, and the release of folded protein to interact with RGS9 and form the functional Gβ_5_-RGS9 dimer. (C) Structures of CCT with different folding substrates: tubulin (left, PDB 7TUB), actin (middle, PDB 7NVM) and Gβ_5_ (right, this study). CCT subunits are labeled and colored based on electrostatic surface charge within the inner wall (red, negatively charged surfaces are labeled; blue, positively charged surfaces).

Cryo-EM image processing revealed an ensemble of snapshots that capture the stepwise Gβ_5_ folding process, beginning with the migration of the unfolded protein to the apical end of CCT2, 5, and 7. The folding process is initiated by specific interactions between the outer β- hairpin of B4 and the inner surface of the CCT5 apical domain, and folding continues radially around the β-propeller as hydrophilic interactions between the outer strands of B3 and B5 with adjacent CCT subunits stabilize their β-sheets. B2 initially adopts an intermediate extended conformation that interacts with the neighboring β-sheet of B3. B6 and B7 then fold without contacting CCT, with their folding presumably stabilized by the other folded blades. The folding of Gβ_5_ is completed with the re-organization of B2 into its native β-sheet and the closing of the β-propeller through the formation of B1. This fully folded Gβ_5_ is presumably then able to bind its RGS binding partner when CCT reopens upon release of the hydrolyzed phosphate.

Our structures reshape the understanding of CCT-dependent protein folding. Based on similarities to the type 1 chaperonin GroEL, previous models have postulated that CCT functions like an “Anfinsen cage” in which an unfolded substrate would bind to the chamber walls of the open form and then release into the space within the CCT folding chamber where it becomes encapsulated in the closed conformation ^44–46^. According to this model, CCT plays a passive role as the substrate folds spontaneously within the protected environment. In contrast, our work reveals that CCT actively directs folding by forming specific interactions with Gβ_5_ to initiate folding and stabilize folding intermediates.

It is notable that folding appears to be initiated between a solvent-exposed β-hairpin of Gβ_5_ with the interior surface of CCT5. Such a minimal cluster of contacts raises the question of how a segment comprising just eight residues (D267 through V274) defines the folding initiation of Gβ_5_. Three distinct contacts are observed at this interface: an electrostatic interaction between R269 (Gβ_5_) and E256 (CCT5), a hydrogen-bond between D267 (Gβ_5_) and the indole nitrogen of W324 (CCT5) and hydrophobic packing between V274 (Gβ_5_) and W324 (CCT5). Interestingly, D267 and R269 are conserved in all five isoforms of Gβ subunits (Gβ1-5), suggesting that these residues play a central role in ensuring proper folding. R269 stands out as the only surface exposed positively charged residue within a large swath of Gβ_5_, and the residue fits precisely in a pronounced negatively charged groove within CCT5 (Figures 3A, B). It is therefore plausible that the β-hairpin containing R269 samples a range of conformations in the molten globule state of Gβ_5_ before its stabilization by E256 of CCT5 and other proximal interactions locks this intermediate in place to enable folding to progress. Consistent with this idea is our observation that the R269E mutation impairs Gβ_5_ folding (Figure 3D, E).

A recent study examining CCT-dependent tubulin folding reported that folding is mediated by electrostatic interactions and hydrogen bonds between CCT and surface residues of tubulin, guiding the folding of individual domains that then consolidate into the native state ^36^. Interestingly, the interactions between CCT and tubulin are distinct from those of Gβ_5_ observed in our study (Figure 7C). Negatively charged surface residues of tubulin associate with the inner walls of the predominantly positively charged CCT hemisphere encompassing CCT1, 3, 6 and 8^36^. With Gβ_5_, the outer blades of the β-propeller also form salt bridges and hydrogen bonds but do so with the inner walls of the predominantly negative charged hemisphere of CCT2, 5 and 7. Together, these findings establish a new paradigm for CCT-dependent protein folding in which CCT in the closed form presents a variety of electrostatic and hydrophilic binding sites that make substrate-specific contacts with surface residues that stabilize folding intermediates and allow the hydrophobic cores to coalesce into the native structure.

The CCT-PhLP1-Gβ_5_ structure and the PhLP1 truncation data rationalize the central role of PhLP1 in Gβ_5_ folding and Gβ_5_-RGS9 dimer formation. In the CCT open form, PhLP1 stabilizes Gβ_5_ binding to CCT, apparently without directly interacting with it. This allosteric effect could occur through binding of the PhLP1 thioredoxin domain to the apical domain of CCT6 in a position that would inhibit CCT closure (Figure 1B), thus trapping Gβ_5_ between the rings in an unfolded state. Evidence supporting this model comes from the intermediate state of CCT in which the high ATPase hemisphere (CCT2, 4, 5 and 7) adopts a closed conformation while the low ATPase hemisphere (CCT1, 3, 6 and 8) remains in an open state and bound to the PhLP1 thioredoxin domain. CCT closure displaces the PhLP1 thioredoxin domain into one folding chamber and creates a positively charged groove between CCT1′ and 4′ into which the negatively charged PhLP1 C-terminal tail binds, tethering the thioredoxin domain in the folding chamber.

The repositioning of PhLP1 upon CCT closure provides insight into its role as a co chaperone. In the closed form, density for PhLP1 becomes well-ordered within CCT, revealing a long α-helix (H1) from the PhLP1 N-terminal domain that traverses between the CCT rings into the opposite folding chamber (Figure 5). This positioning of PhLP1 in the closed conformation would clash with Gβ_5_ bound between the CCT rings in the open state. Thus, PhLP1 may help displace Gβ_5_ from between the rings as the chamber closes. Moreover, the PhLP1 N-terminal domain extends across the folding chamber containing Gβ_5_ to the positively charged groove at the CCT1/4 interface where a sequence of three phosphorylated serine residues on PhLP1 (S18-20) bind. Notably, previous work has shown that phosphorylation of these sites is required for efficient Gβγ assembly ^15, 16^, and our results extend this observation to Gβ_5_-RGS9 assembly (Figures 6C, D). The structure suggests that this interaction could assist in Gβ_5_ folding in two ways. First, insertion of acidic motifs into the positively charged grooves between CCT1 and CCT4 at both ends would counteract the potential charge repulsion between the two subunits as CCT transitions between the open and closed states (Figure 5C). Second, the location of phosphorylated S18-20 in the positively charged groove requires that the PhLP1 N-terminal extension traverses the folding chamber, possibly wrapping around Gβ_5_ and constraining it to the hemisphere encompassing CCT2, 4, 5 and 7 where folding of the β-propeller occurs.

Visualizing protein folding has been a major challenge in structural biology ^47^. However, recent advances in cryo-EM image processing that enable structural determination in a continuous array of functional states have allowed us to elucidate the CCT-mediated folding trajectory of the G protein β_5_ subunit. This work establishes a structural framework to investigate the effects of pathogenic mutations in Gβ_5_ that are associated with protein misfolding^48–50^ and to determine the folding trajectories of other β-propeller proteins, including the other Gβ subunits, the mLST8 and Raptor subunits of mTOR complexes, and the BBS7 subunit of the BBSome complex, among others ^20, 39, 51^. With various CCT substrates now known to bind to different CCT subunits in their closed state, it appears likely that substrates have co-evolved with CCT to preserve optimal binding sites to initiate folding.

## Acknowledgements

This work was supported by grants to P.S.S. (NIH R35 GM133772), B.M.W. (NIH R01 EY012287), and a fellowship from the Brigham Young University Simmons Center for Cancer Research to Y.K. We thank the University of Utah Arnold and Mabel Beckman Center for Cryo-EM and Center for High Performance Computing for cryo-EM and computational support, respectively. We also thank Daniel S. Bradford and Samuel L. Cottam for technical assistance.

## Author Contributions

Conceptualization, B.M.W. and P.S.S.; Methodology, B.M.W., P.S.S., S.W., M.I.S, and Y.K.; Investigation, S.W., M.I.S., Y.K., W.G.W., T.M.S., E.J.C., N.E.G., and P.S.S.; Data Curation, S.W. and P.S.S.; Writing – Original Draft, B.M.W. and P.S.S., Writing – Review & Editing, B.M.W. and P.S.S.; Visualization, S.W., M.I.S., Y.K., W.G.W., M.R, J.H.I, B.M.W., and P.S.S.; Funding Acquisition, B.M.W. and P.S.S.; Supervision, B.M.W. and P.S.S.

## Methods

### Cell culture

HEK-293T cells were cultured in DMEM/F-12 supplement with 10% FBS in T-25 flasks. At 40-50% confluency, cells were transfected with 540 µl of transfection reagent containing 2 µg of an N-terminal 2x strep peptide-tagged human Gβ_5_ cDNA or Gβ_5_ R269E construct in the pcDNA3.1 vector and 2 µg of a C-terminal myc-His tagged human wild type PhLP1, PhLP1 truncation, or PhLP1 S18-20A cDNAs in the pcDNA3.1 vector along with 2 μg of a C-terminal HA tagged human RGS9 cDNAs in the pcDNA3.1 vector as indicated. Control transfections contained variable amounts of GFP in the same vector to make a total of 8 μg cDNA added.

### Immunoblot

Cells were washed in phosphate-buffered saline (PBS) and lysed in 500 µl of lysis buffer (PBS supplemented with 1% IGEPAL, 0.5 mM PMSF and Halt Protease Inhibitor Cocktail).

Lysates were triturated 10 times through a 25-gauge 7/8” needle, then centrifuged at 21,000 x g for 10 minutes at 4°C to clarify the lysates. CCT was immunoprecipitated from the lysate by adding 4 µg of a CCT5 antibody (BioRAD MCA2178, 1:100 IP) and incubating for 30 minutes at 4°C, followed by adding 25 µl of protein A/G agarose beads for another 30 minutes at 4°C. RGS9 was immunoprecipitated from the lysate by adding 1 μg of a HA antibody (Sigma Aldrich 1186723001, 1:400 IP) following the same protocol. Immunoprecipitates were washed three times in lysis buffer, then resuspended in 30 µl of SDS-PAGE sample loading buffer. Proteins were separated by SDS-PAGE on 10% gels and transferred to nitrocellulose. The nitrocellulose was probed for 16 hrs at 4°C with the indicated primary antibodies at the following dilutions in TBS blocking buffer (LI-COR 927-60001): Strep (Genscript A01732, 1: 5,000), c-myc (Invitrogen 13-2500, 1: 1,000), CCT2 (Abcam, Ab92746, 1: 10,000), HA (Sigma Aldrich 1186723001, 1:1000), and GFP (Abcam Ab6556, 1: 5,000). Blots were wash 3 times with TBS-Tween wash buffer (50 mM TrisHCl pH 7.6, 150 mM NaCl, 50 mM NaOH, 0.05% Tween 20) and probed with appropriate IRDye-labeled secondary antibodies at 1:10,000 dilutions in TBS Tween wash buffer: IRDye 680RD Goat Anti-Mouse (LI-COR 926-68070), IRDye 800RD Goat Anti-Rabbit (LI-COR 926-32211), and IRDye 800CW Goat Anti-Rat (LI-COR 926-32219).

Blots were imaged using a LI-COR Odyssey infrared scanner, and band intensities were quantified with the LI-COR ImageQuant software.

### Purification of CCT-PhLP1-Gβ_5_

HEK-293T cells were cultured in DMEM/F-12 supplement with 10% FBS in T-25 flasks. At 40% confluency, cells were transduced with pLenti-Puro virus containing human Gβ_5_ with N-terminal 2X Strep and HPC4 tags. The cells were then treated with puromycin for one week to select for transduced cells. The selected cells were then transduced with pLenti-Blast virus containing human PhLP1 with N-terminal His_6_ and Myc tags. The cells were then treated blasticidin for another week to select for transduced cells. The selected cell line was frozen in aliquots and used as a stable cell line for CCT-PhLP1-Gβ_5_ purification.

HEK-293T cells stably expressing Strep-Gβ_5_ and His-PhLP1 were grown to 80% confluency in T-175 tissue culture flasks, harvested and lysed in lysis buffer at 5 mL of buffer per gram of harvested cell pellet. The lysate was centrifuged at 30,000 x g for 20 minutes and filtered through a 0.45 μm filter followed by a 0.22 μm filter. The CCT-PhLP1-Gβ_5_ complexes were then purified at 4° C using a tandem affinity approach. The filtered lysate was passed over a HisTrap HP 5 mL column equilibrated with 20 mM HEPES, 20 mM NaCl, 25 mM imidazole, 0.05% CHAPS, 1 mM TCEP, pH 7.5 for an hour. The column was then washed with 5 column volumes of equilibration buffer, and protein was eluted with linear gradient of 8 column volumes of 25 mM to 500 mM imidazole. Eluted fractions were analyzed by SDS-PAGE and Coomassie staining. Fractions containing PhLP1-CCT were combined and loaded onto 3 mL of Strep-Tactin resin equilibrated with 20 mM HEPES pH 7.5, 20 mM NaCl for an hour. The column was washed twice with one column volume of 20 mM HEPES pH 7.5, 150 mM NaCl, and then twice with one column volume of 20 mM HEPES pH 7.5, 20 mM NaCl. The CCT-PhLP1-Gβ_5_ complexes were eluted with 3 column volumes of 20 mM HEPES pH 7.5, 20 mM NaCl, 0.05% CHAPS, 10 mM D-desthiobiotin, 1 mM TCEP, and 5 mM MgCl_2_ and concentrated to 1.5 μg/μL using a 30 kDa cutoff filter and analyzed by SDS-PAGE and Coomassie staining and immunoblotting.

Endogenously expressed human CCT complex without Gβ_5_ or PhLP1 was purified from HEK-293T cells as described previously ^39^ and was also analyzed by SDS-PAGE and Coomassie staining and immunoblotting. To assess the functional state of this PhLP1-free CCT and the CCT-PhLP1-Gβ_5_ complex, their ATPase activity was measured using the Malachite Green Phosphate Assay Kit (Sigma-Aldrich MAK307). This assay measures inorganic phosphate released upon ATP hydrolysis. ATP at 0.25 mM final concentration was added to freshly prepared CCT complexes (0.5 μM final concentration) in 20 mM HEPES, 20 mM NaCl, 5 mM MgCl_2_, pH 7.5 in a total volume of 80 μl. ATP hydrolysis was allowed to proceed for 15 min. at 23°C, after which 20 μl of the Malachite Green reagent was added and incubated for an additional 15 min. at 23°C and the absorbance was measured at 640 nm.

### Chemical crosslinking coupled with mass spectrometry

Purified CCT-Gβ_5_-PhLP1 complex (50 to 150 μg) was diluted to 1 mg/mL in crosslinking buffer (20 mM HEPES, 20 mM NaCl, 5 mM MgCl_2_, pH 7.5). NHS-Long chain-diazirine (LC-SDA, 150 μg) was dissolved in 6 μL acetonitrile and 144 μL crosslinking buffer to make a 1 mg/mL solution. The LC-SDA was added to the purified complex at 0.22 μg LC-SDA for every 1 μg of protein and the mixture was incubated in the dark at room temperature for 1 hr. The reaction was quenched with 50 mM TrisHCl. The sample was then loaded onto the lid of an Eppendorf tube and incubated on ice under UV light (365 nm source) 9 cm from the source for 30 min at 200 mJ/cm^3^. The crosslinked sample was denatured with 100 μL of 6M guanidine and then concentrated on a 10 kDa cutoff filter to a volume of 100 μL to remove peptide fragments. The sample was alkylated by adding iodoacetamide to 10 mM and incubated 15 min in the dark at room temperature. The sample was reduced by incubating for 30 min at 37 °C with 5 mM TCEP. Ammonium bicarbonate (400 μL of 500 mM) was added to quench the reaction. The sample was then digested overnight at 37°C with 0.5 μg of trypsin. After 16 hr, the digestion was stopped with 10% trifluoroacetic acid, and the sample was desalted using C18 stage tips equilibrated with 5% trifluoracetic acid. The sample was run over the column 5 times and washed twice with 100 μL of 5% trifluoroacetic acid. To elute, 50 μL of 70% acetonitrile was passed over the column 3 times, allowing the second elution volume to sit on the column for 1 min before eluting. The eluted protein was collected in an Eppendorf tube and vacuum concentrated to dryness. Sample was resuspended in 50 μL of AspN digestion buffer (50 mM TrisHCl, 0.5 mM zinc nitrate, pH 8.0), and 0.5 μg of AspN was added to the sample and incubated overnight at 37 °C. After 16 hr, the digestion was quenched with 10% trifluoroacetic acid. To enrich for crosslinked peptides, the sample was loaded onto a size exclusion column equilibrated with 2 column volumes of 30% acetonitrile and 0.1% trifluoroacetic acid in water.

Sample was eluted with 1 column volume of 30% acetonitrile and 0.1% trifluoroacetic acid in water at a flow rate of 0.05 mL/min, collecting fractions every 100 μL. The first two high molecular weight fractions, composing the first half of the elution peak, were collected and desalted using C18 stage tips as previously described. The sample was vacuum concentrated to dryness and resuspended in 3% acetonitrile and 0.1% formic acid in water.

Samples were analyzed by LC-MS using an EASY-nlc 1200 LC system coupled to a Orbitrap Fusion Lumos Tribrid mass spectrometer equipped with an EASY Spray source. Mobile phase A consisted of 3% acetonitrile and 0.1% formic acid in water and mobile phase B consisted of 80% acetonitrile and 0.1% formic acid in water. Peptides were loaded into an EASY-Spray column operated at a temperature of 35°C and a flow rate of 0.3 μL/min. Peptides were separated at a flow rate of 0.3 μL/min using the following gradient: 5% mobile phase B (0-2 minutes), 2-32% mobile phase B (2-87 min), 32-42% mobile phase B (87-102 min), 42-95% mobile phase B (102-112 min), 95% mobile phase B (112-122 min), 95-2% mobile phase B (122-125 min), 2% mobile phase B (125-127 min), 2-100% mobile phase B (127-129 min), 100% mobile phase B (129-132), 100-2% mobile phase B (132-133 min), 2% mobile phase B (133-135 min), 2-100% mobile phase B (135-137 min), 100% mobile phase B (137-140 min), 100-0% mobile phase B (140-142 min), 0% mobile phase B (144-150 min).

MS data were acquired using the Orbitrap to detect both MS and MS/MS spectral scans. The data was collected in a data-dependent mode with a cycle time of 3 s. MS1 spectra were acquired at a resolution of 120,000 using a scan range of 375-1,700 m/z and AGC target of 4.0 x 10^5^ with a maximum injection time of 50 s. Monoisotopic peak determination (MIPS) was activated, and only precursors with charge states between 2 and 6 with an intensity higher than 5,000 were selected for fragmentation. Selected precursors were fragmented by CID using a collision energy setting of 30%. MS2 spectra were acquired at an AGC of 10,000 and a maximum injection time of 35 ms. Dynamic exclusion was set to 60 s after 1 count.

The crosslink search was performed using xiSEARCH v1.7.6.7 ^52^ with the following parameters: MS accuracy 6 ppm; MS/MS accuracy 20 ppm; enzymes trypsin and AspN; exclusion of non-covalent interactions activated; missed cleavages allowed 4.

Carbamidomethylation of cysteine residues was set as a fixed modification, while the following were set as variable modifications: oxidation of methionine, LC-SDA crosslinker alone (mass modification: 223.13207784 Da), hydrolyzed LC-SDA (mass modification: 213.136494 Da), LC-SDA alkylation (mass modification: 181.110279 Da), and LC-SDA oxidation (mass modification: 211.120839 Da). LC-SDA specificity for lysine, serine, threonine, and tyrosine residues, and N-terminus for the NHS reaction, and any residues for the diazirine reaction.

Spectra were searched against FASTA files containing the human sequences for each of the eight CCT subunits, Strep peptide and HPC4 tagged Gβ_5_, and His_6_ and Myc tagged PhLP1. A FASTA file containing the reverse sequences of the target proteins was also included as a decoy database. FDR was estimated at a 5% residue pair level using xiFDR v2.1.5.5, including only unique peptide spectra match with a minimum peptide length of 5 residues, excluding consecutive peptide links, and boosting residue pairs between different proteins.

As an internal control of the reliability of the dataset, crosslink Cα distances between residues were mapped within the equatorial domains, the most structurally stable region of CCT. The maximal possible crosslinking distance for LC-SDA is 30-40 Å, which takes into account the crosslinker arm (12.5 Å), the side chain lengths (variable up to 12.6 Å) plus any protein backbone dynamics around the two crosslink sites. Greater than 80 % of the crosslink distances were less than 40Å, indicating a good quality dataset.

### Inducing the closed conformation of CCT

To 10 µl of 1.5 mg/ml co-purified CCT-PhLP1-Gβ_5_, 0.5 μL of 600 mM KCl, and 1.5 µl each of 10 mM Al(NO_3_)_3_, 60 mM NaF, and 10 mM ATP were sequentially added in rapid succession to avoid Al(OH)_3_ precipitation. The complex was then incubated at 37°C for an hour and centrifuged at 16,300 x g for 1 min to remove any precipitate.

### Electron cryo-microscopy specimen preparation

UltrAuFoil R2/2 Au200 mesh grids (Quantifoil) were glow discharged for 1 min on both sides at 25 mA using a Pelco easiGlow unit (Ted Pella, Inc.). 3 µL of freshly prepared CCT-PhLP1-Gβ_5_ sample treated with ATP and AlF_x_ were applied on the grids and blotted with filter paper (595 Filter Paper, Ted Pella, Inc.) for 2.5 s using a Mk. II Vitrobot (Thermo Fisher Scientific) with a wait time of 25 s and -1 mm offset. The grid was then plunge frozen into liquid ethane and stored in liquid nitrogen until cryo-EM imaging.

Cryo-EM movies were recorded using SerialEM v3.8 ^53^ in super-resolution mode on a 300 kV Titan Krios (Thermo Fisher Scientific) equipped with a post-GIF K3 direct detector (Gatan, Inc.). A total of 24,986 movies were recorded at a nominal magnification of 81,000X, corresponding to a super-resolution pixel size of 0.529 Å with a total dose of 40.42 e/Å^2^ and 40 frames per movie. Cryo-EM data collection parameters are summarized in Table 2.1.

### Image processing

Super-resolution cryo-EM movie frames were motion corrected, dose weighted, Fourier binned 2X, and summed using cryoSPARC Live as implemented in cryoSPARC v3.3 ^54^. CTF parameters were determined using patch CTF estimation. The total exposures were then curated using CTF fit resolution cutoff of 5 Å.

A total of 6,077,238 particles were selected across 19,137 curated micrographs using a blob picker with a 180-220 Å particle diameter restraint. Particles were then extracted with a box size of 300 pixels and Fourier cropped to 150 pixels (2x binned). Four rounds of 2D classifications were performed, and a total of 2,373,004 selected particles were then used for six classes of heterogeneous reconstructions with input volume of opened, intermediate, and closed CCT derived from ab initio reconstructions. A total of 1,895,426 particles were sorted into the closed state, and 279,293 particles were sorted into the open state.

Further heterogeneous refinement on open particles revealed that 137,857 particles were in a canonical open state and 114,711 particles were in an intermediate, partially-closed state with resolution at 3.4Å and 3.7Å, respectively. Heterogeneous refinement followed by non-uniform refinement was then performed on open particles, which result in 104,907 particles in the open state with 3.4Å resolution.

Closed particles were subjected to another round of heterogeneous refinement to remove junk particles. A total of 1,813,701 closed state particles were sorted into well-resolved 3D reconstructions and re-extracted to the unbinned 300-pixel box size. Three modes of focused 3D variability analysis (3DVA) with 6 Å filtered resolution and a mask covering the substrate density were performed. Initial analysis for 3DVA found that the Gβ_5_ density in component 0 had significant continuous changes. 5 clusters of individual intermediate states were then resolved from component 0 using 3DVA display with 6Å filtered resolution and followed by non-uniform refinement. Each cluster was further processed using focused 3D classification without realignment of particle orientations using a mask that only covers the Gβ_5_ density.

Focused 3D classification was performed with five classes, and 6Å target resolution and PCA initialization mode. Five classes from each cluster were resolved and subjected to non-uniform refinement.

A total of 25 classes with a resolution range between 2.8-3.0 Å were generated from focused 3D classification. Of these, 16 distinct classes were resolved with different Gβ_5_ folding intermediates. These classes were ordered based on increasing completeness of Gβ_5_ density (classes 0-15).

The next component, component 1, from 3DVA showed significant heterogeneity within PhLP1 density. Further 3DVA display using this component resolved five clusters with various intensities of PhLP1 densities. The class with best resolved PhLP1 density was selected for non uniform refinement for further resolution improvement. A detailed image processing workflow is presented in Extended Data Figure 8.

### Model building and refinement

The models of Gβ_5_ and PhLP1 in closed CCT were refined from AlphaFold Gβ_5_ model (AF-14775-F1) and AlphaFold PhLP1 model (AF-Q1337-F1). The initial model was docked into each intermediate state map as a rigid body using UCSF Chimera (version 1.15) ^54^. A detailed model was manually rebuilt and refined in COOT v0.9.6 ^55^. The PhLP1 model was further refined using real-space refinement in Phenix v1.19.2 ^56^. Gβ_5_ was further refined with its neighboring CCT subunits (PDB:7nvm ^37^), CCT 2, 4, 5 and 7, using real-space refinement in Phenix v1.19.2. The rest of the CCT subunits were added into the refined PhLP1 and Gβ_5_ model using Chimera and combined into a single CCT-PhLP1-Gβ_5_ model. The complex model was then subjected to real-space refinement in Phenix with a supplement of ADP, AF3, HOH, Mg^2+^ restraint .cif files. Model refinement statistics are summarized in Table 2.2.

### AlphaFold prediction

ColabFold as implemented in AlphaFold2 was used to predict the interaction between CCT1 and the PhLP1 negatively charged sequence near the N-terminus. The input sequence for CCT1 was CCT1 A178-N217, L312-S405, and the input sequence for PhLP1 N-terminal domain was S18-D29 where S18-S20 were changed to glutamates to as phosphomimetics of the phosphorylated serine residues. The final input sequence was: AIKYTDIRGQPRYPVNSVNILKAHGRSQMESMLISGYALN: LKRDLKRIAKASGATILSTLANLEGEETFEAAMLGQAEEVVQERICDDELILIKNTKARTSASIILRGANDFMCDEMERSLHDALCVVKRVLES:EEEEDEDSDHED. The template_mode was set to pdb70. The num_recycles was set to 24.

