## Supplemental Figures for "Visualizing the chaperone-mediated folding trajectory of the G protein β5 β-propeller"

**Supplemental Material**

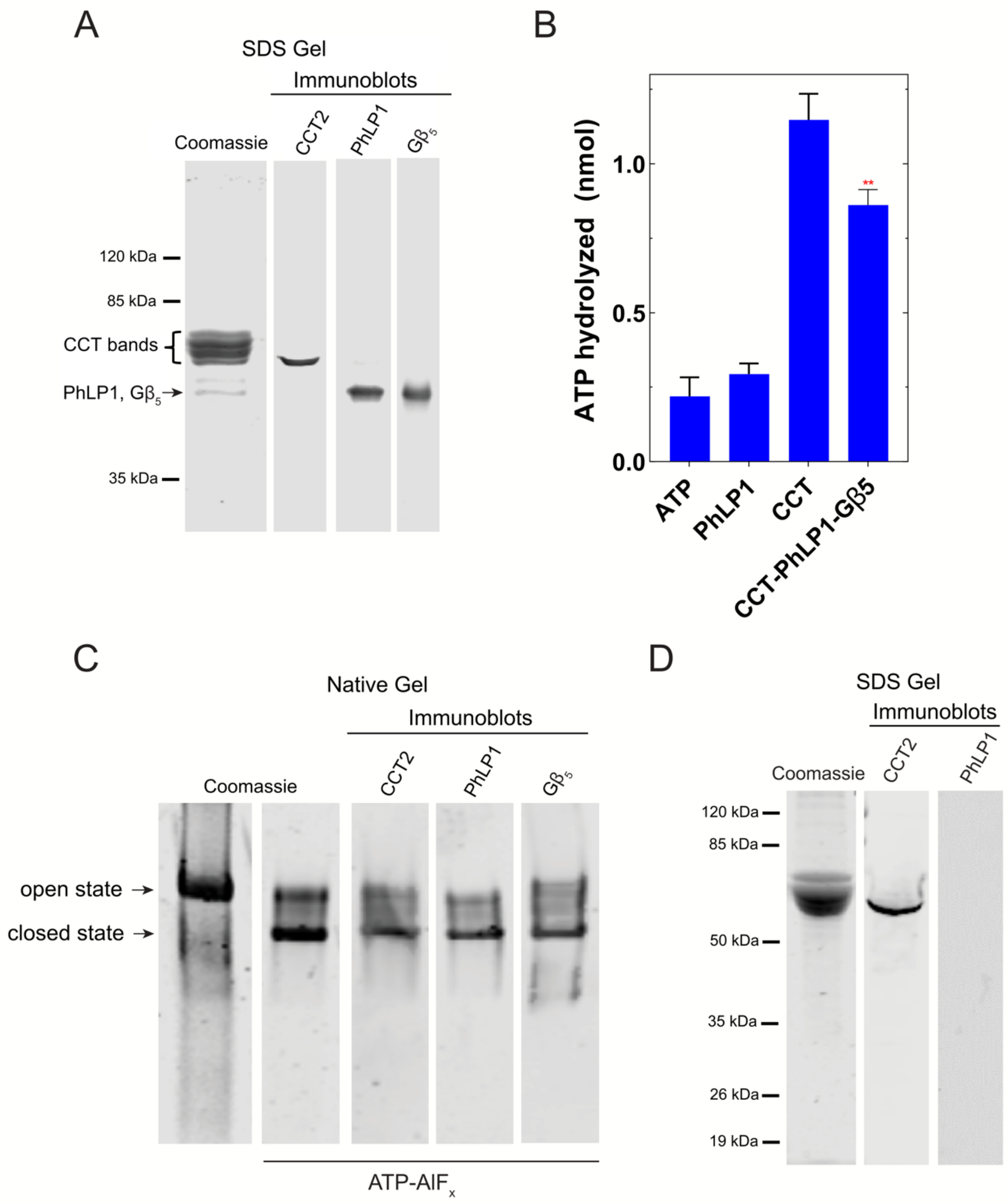


**Supplemental Figure S1. Characterization of purified CCT complexes.** (A) Coomassie-stained denaturing gel of purified human CCT-PhLP1-G$\beta_5$ and immunoblots for CCT2, PhLP1 and G$\beta_5$. (B) ATPase activity of the CCT-PhLP1-Gβ5 complex compared to CCT alone. Red ** $p < 0.01$ in a t-test compared to CCT. (C) Coomassie-stained native PAGE and immunoblots of purified CCT-PhLP1-G$\beta_5$ with and without ATP-$AlF_x$. Positions of the opened and closed state CCT are indicated. (D) Coomassie-stained denaturing gel and immunoblots of purified human CCT in the absence of PhLP1.

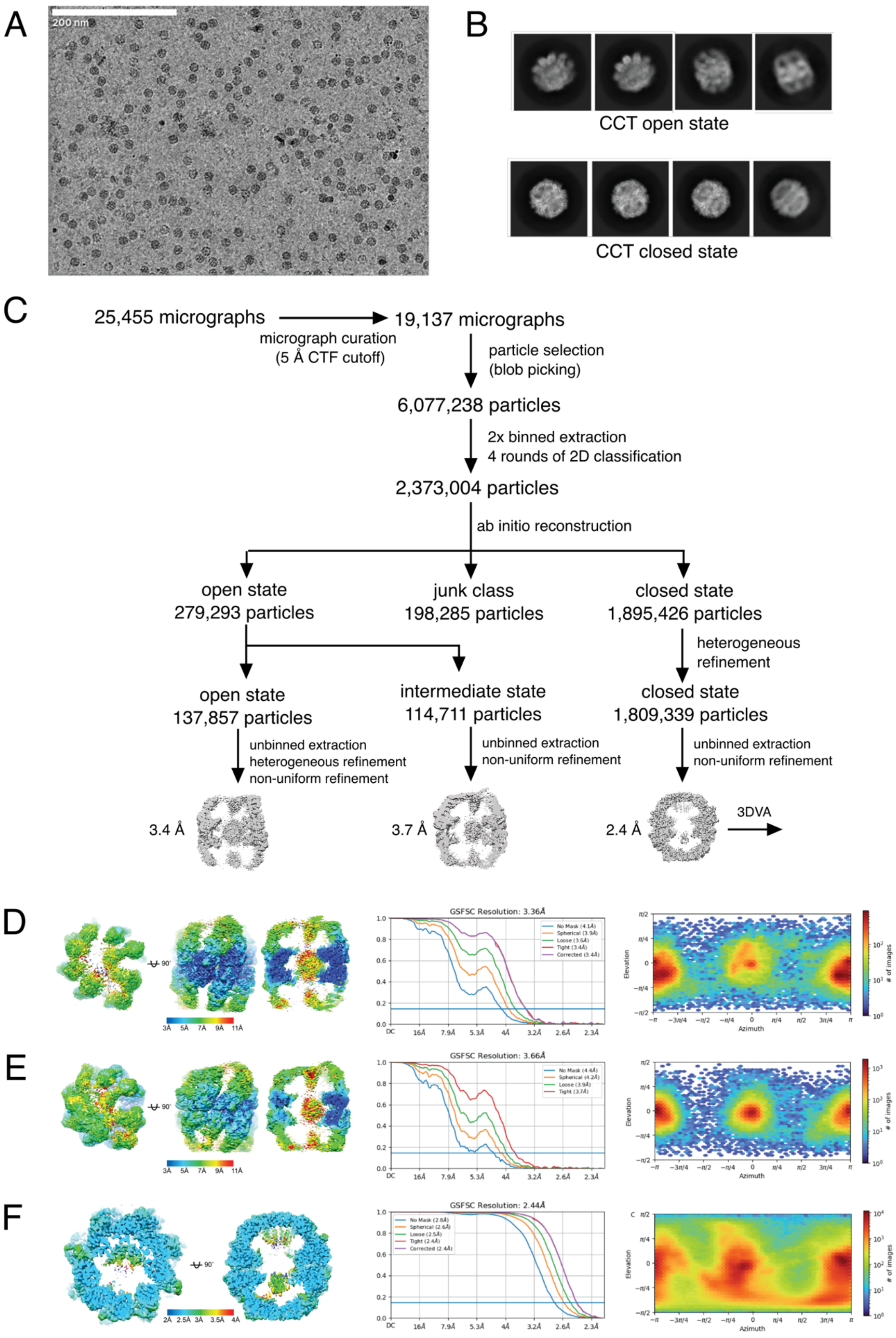


**Supplemental Figure S2. Image processing workflow of G$\beta_5$-PhLP1-CCT particles.** (A) Representative cryo-EM micrograph, (B) reference-free 2D class averages, and (C) processing workflow for open, intermediate, and closed state of CCT. (D-F) Cryo-EM validation of open (D), intermediate (E), and closed (F) reconstructions. Left, Reconstruction heat maps; middle, gold-standard FSC plots; right, particle orientation distribution.

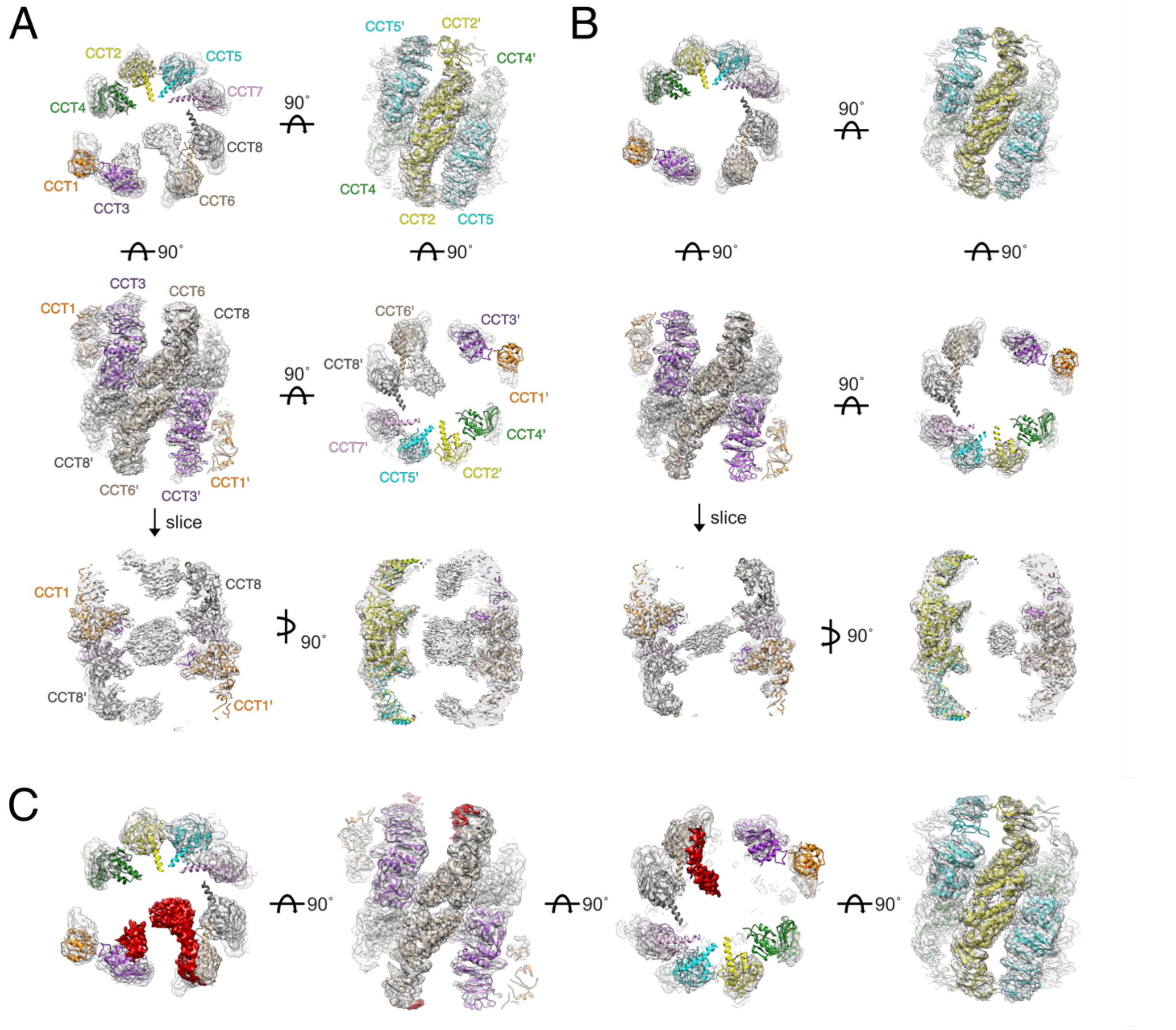


**Supplemental Figure S3. Comparison of CCT reconstructions with and without PhLP1.** (A) Reconstruction of open state CCT with unfolded G$\beta_5$ and PhLP1 fitted with a model of CCT (PDB 6QB8). (B) Reconstruction of CCT without PhLP1. (C) Difference map of reconstructions with and without PhLP1 reveal PhLP1-specific densities (red). The difference map is superimposed on the reconstruction of CCT without PhLP1 (gray).

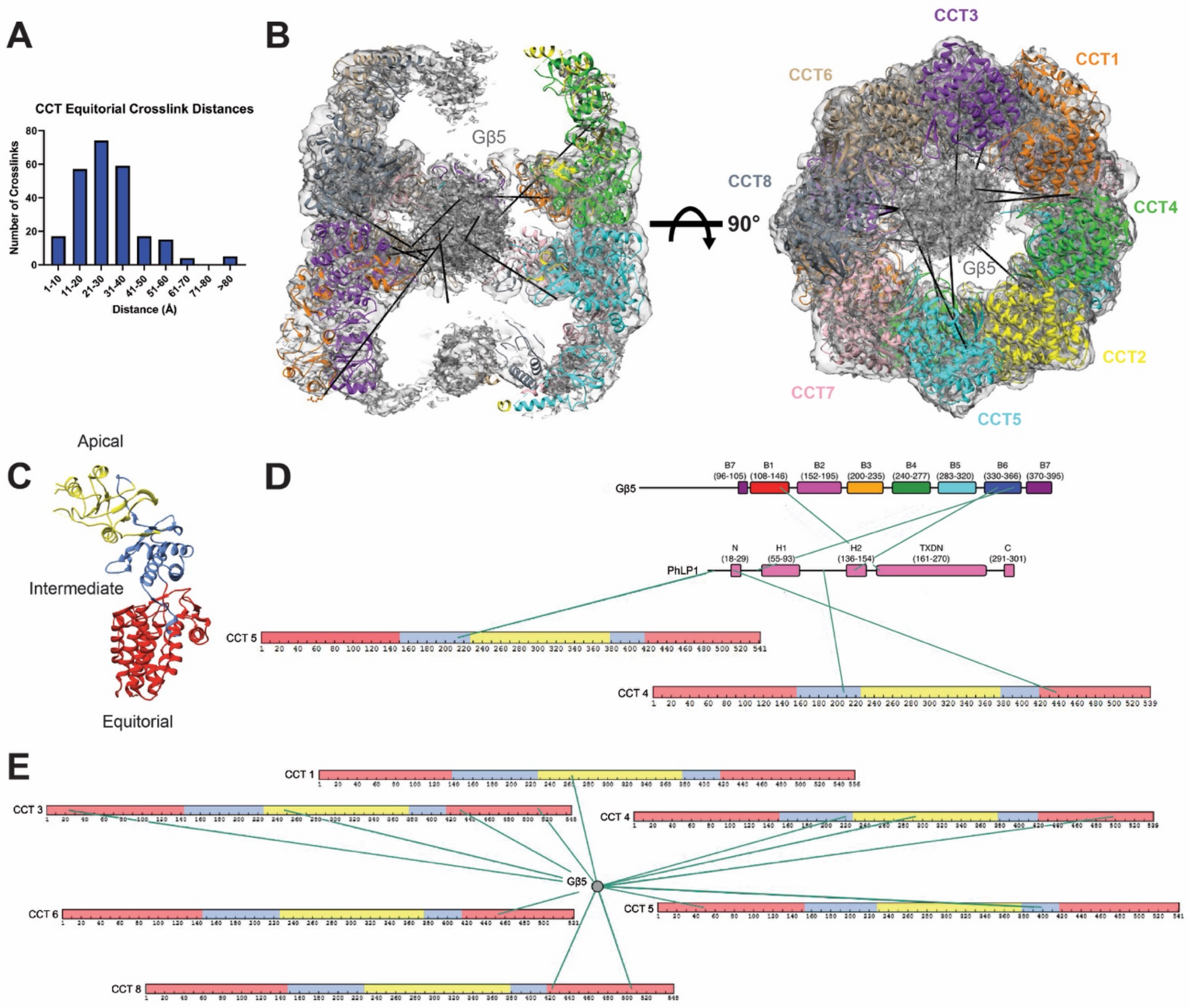


**Supplemental Figure S4. Crosslinking of the CCT-PhLP1-G$\beta_5$ complex in the open conformation.** (A) Quantification of crosslink distances within the CCT equatorial domains. Greater than 80% of the crosslinks were < 40 Å, indicating a quality dataset. (B) Crosslinks of unstructured G$\beta_5$ to CCT subunits from a cutaway side view and top view. (C) A model of the domains of a CCT subunit. The coloring corresponds to the sequences in D and E. (D) PhLP1 crosslinks to G$\beta_5$ and CCT subunits modeled onto a sequence map. Only unique crosslinks are shown. (E) G$\beta_5$ crosslinks to CCT subunits modeled onto a sequence map. Only unique crosslinks are shown. The G$\beta_5$ sequence is not modeled as G$\beta_5$ is unstructured in the open form of CCT.

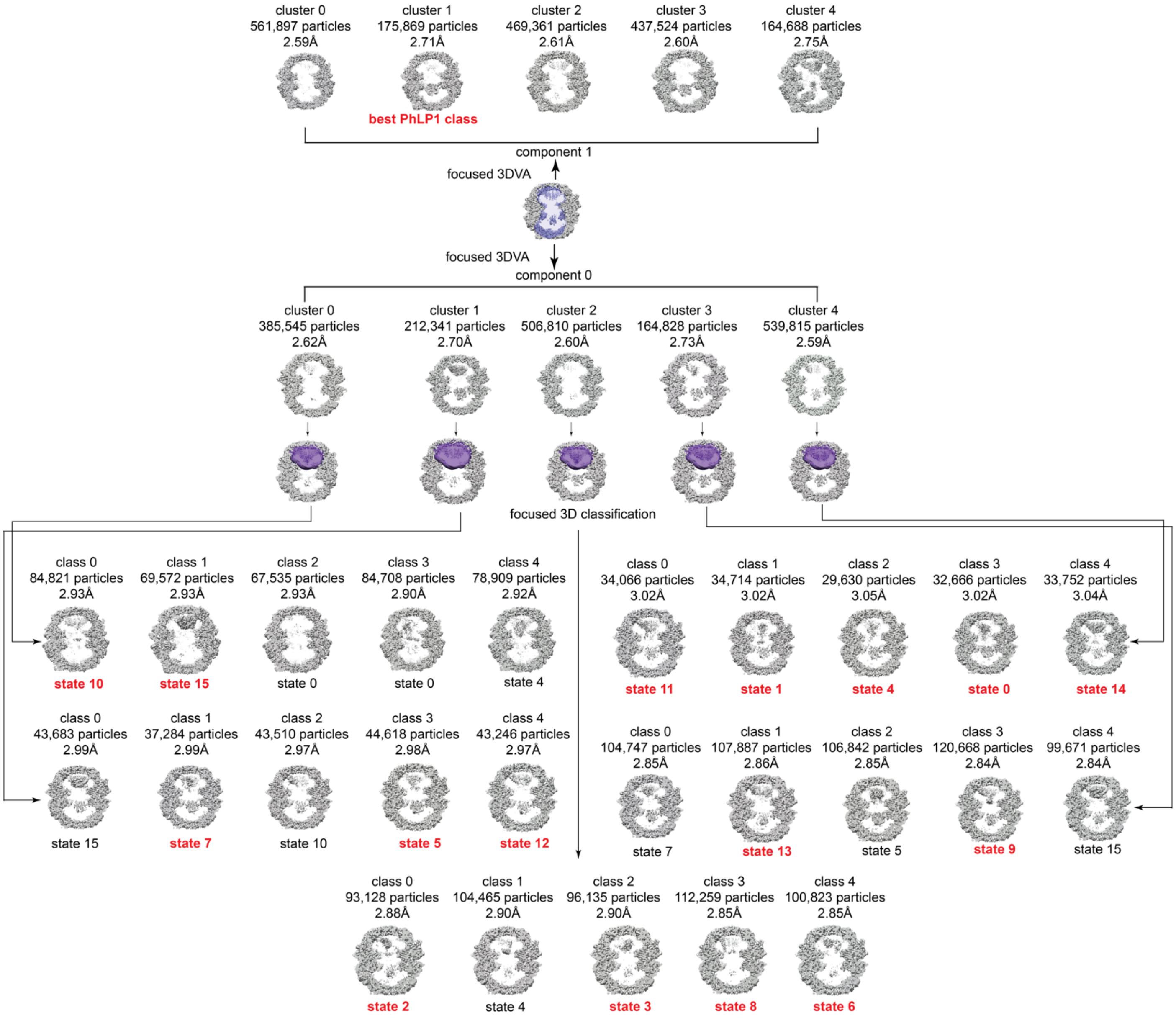


**Supplemental Figure S5. Workflow for 3D variability analysis.** The mask used in focused classification is indicated in purple. Maps that were selected for model building are highlighted in red (based on the state with the highest local resolution of PhLP1 or G$\beta_5$).

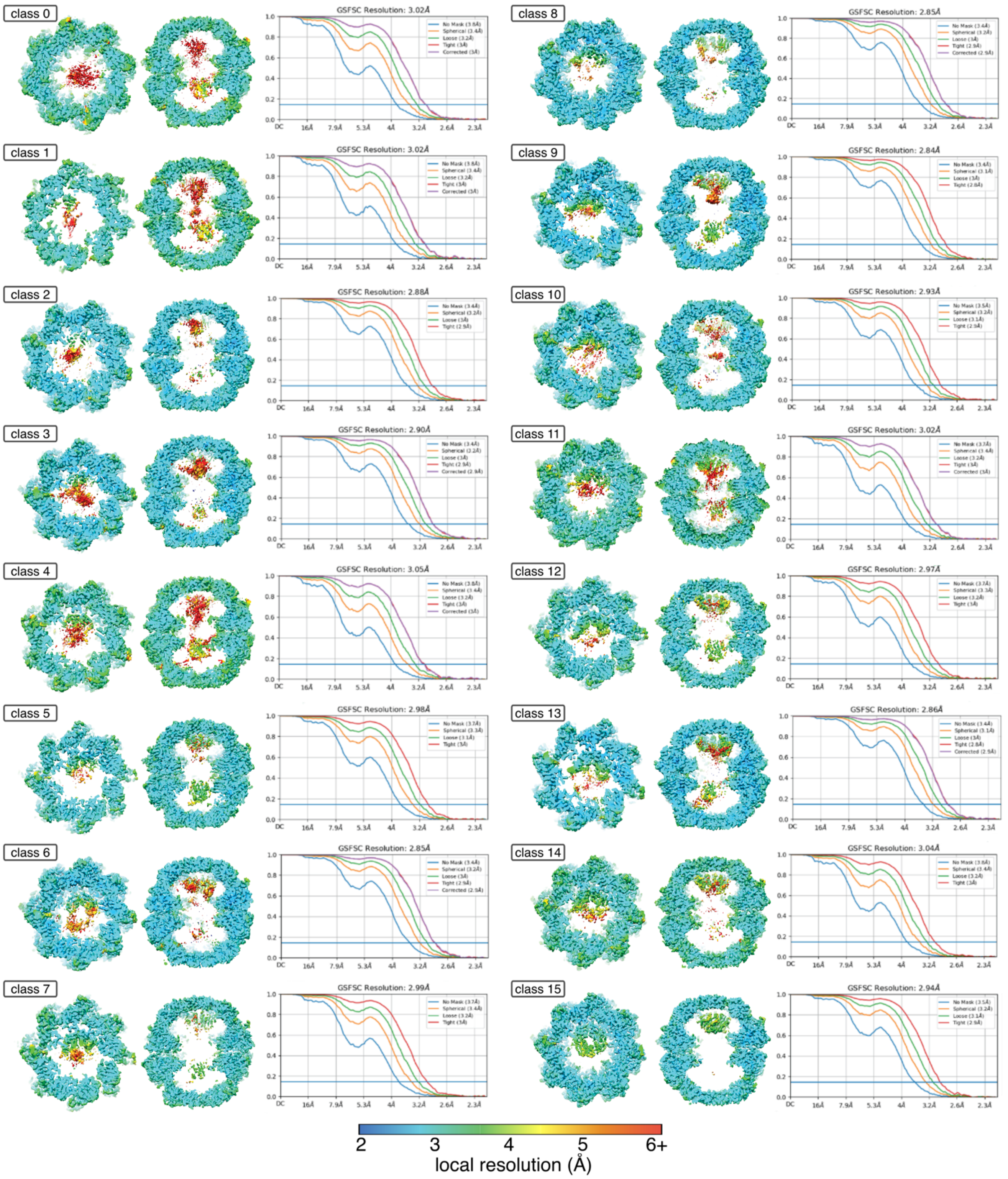


**Supplemental Figure S6. Reconstructions of closed states 0-16.** Local resolution heat maps (top and side sliced views) and gold standard FSCs for each class are shown.

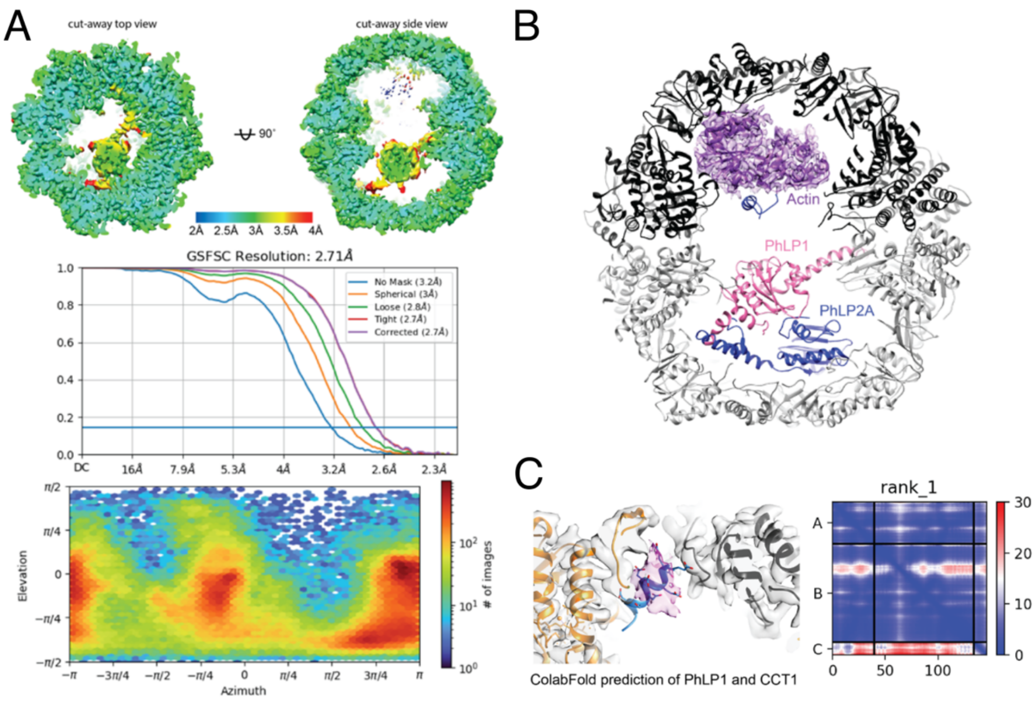


**Supplemental Figure S7. PhLP1 structural analysis.** (A) Cryo-EM validation of the 3D reconstruction used for PhLP1 modeling, including local resolution heat map (top), gold standard FSC (middle), and orientation distribution (bottom). (B) Overlay of PhLP1 (this study) and PhLP2A (PDB 7NVM [37]) models bound to closed CCT. Actin model and density shown as a reference for the PhLP2A structure. (C) Left, ColabFold prediction of the position of PhLP1 residues 18-29 binding to CCT1. Putative PhLP1 density is shown in purple. Serines 18-20 were changed to glutamate to mimic phosphorylated serines. Right, Predicted aligned error (PAE) of the CCT1-PhLP1 ColabFold prediction. Chain A represents CCT1 L312-S405, Chain B represents CCT1 A178-N217, Chain C represents PhLP1 S18-D29 with the S18-S20E change. Distance errors ranges from 0 (blue) to 30 Å (red).

### Multimedia Files

**Supplemental Movie 1.** Snapshots of classes 0-16 assembled into a "flipbook" of G$\beta_5$ folding. The left panel shows the local resolution of G$\beta_5$ and surrounding CCT density and the model built into well resolved densities. The right panel shows the color-coded blades according to the figure scheme used throughout the paper.

**Supplemental Movie 2.** Animation of G$\beta_5$ folding by CCT and PhLP1.

**Supplemental Table S1. Cryo-EM Data Collection Parameters**

| **Data collection** | |
|---|---|
| Microscope | Titan Krios G3 |
| Voltage (kV) | 300 |
| Detector | Gatan K3 |
| Data collection software | SerialEM |
| Nominal magnification | 81,000x |
| Dose rate ($e^-/Å^2$/second) | 16.1 |
| Total number of frames | 40 |
| Total electron exposure ($e^-/Å^2$) | 40.42 |
| Defocus range (μm) | -0.8 to -1.5 |
| Pixel size (Å) | 0.529 (super-resolution) |
| **Data processing** | |
| Number of micrographs | 25,455 |
| Initial particle picked | 6,077,238 |

**Supplemental Table S2. 3D reconstruction and model refinement statistics (1/3)**

| **Structure** | closed state 0 | closed state 1 | closed state 2 | closed state 3 | closed state 4 | closed state 5 |
|---|---|---|---|---|---|---|
| EM Databank Accession ID | EMD-40440 | EMD-40452 | EMD-40454 | EMD-40453 | EMD-40482 | EMD-40488 |
| Protein Data Bank Accession ID | 8SFF | 8S8G | 8SGC | 8SG9 | 8SHA | 8SHL |
| **Symmetry imposed** | | | | | | |
| 3D classification | C1 | C1 | C1 | C1 | C1 | C1 |
| 3D refinement | C1 | C1 | C1 | C1 | C1 | C1 |
| Final particles | 32,666 | 34,714 | 93,128 | 96,135 | 29,630 | 44,618 |
| **Map resolution (Å)** | | | | | | |
| FSC 0.143 (unmasked) | 3.8 | 3.8 | 3.4 | 3.4 | 3.8 | 3.7 |
| FSC 0.143 (masked, corrected) | 3.0 | 3.0 | 2.9 | 2.9 | 3.0 | 3.0 |
| **Model Refinement** | | | | | | |
| Initial model used for CCT (PDB code) | 7NVM | | | | | |
| Initial model used for G$\beta_5$ (AlphaFold protein structure database) | N/A | AF-O14775-F1 | | | | |
| Initial model used for PhLP1 (AlphaFold protein structure database) | N/A | AF-Q13371-F1 | | | | |
| Map correlation coefficient | 0.88 | 0.870 | 0.880 | 0.880 | 0.840 | 0.880 |
| **Model composition** | | | | | | |
| Non-hydrogen atoms | 66,750 | 66,794 | 66,712 | 65,583 | 67,158 | 67,244 |
| Protein residues | 8,667 | 8,672 | 8,665 | 8,522 | 8,717 | 8,731 |
| Ligands (ADP, $AlF_x$, $Mg^{2+}$) | 16 | 16 | 16 | 16 | 16 | 16 |
| **R.m.s. deviations** | | | | | | |
| Bond lengths (Å) | 0.003 | 0.002 | 0.004 | 0.003 | 0.006 | 0.002 |
| Bond angles (˚) | 0.535 | 0.517 | 0.540 | 0.523 | 0.609 | 0.510 |
| **Validation** | | | | | | |
| MolProbity score | 1.590 | 1.590 | 1.620 | 1.560 | 1.69 | 1.550 |
| Clashscore | 7.12 | 7.140 | 7.060 | 6.670 | 7.29 | 6.870 |
| Poor rotamers (%) | 0.00 | 0.000 | 0.000 | 0.000 | 0.000 | 0.000 |
| **Ramachandran plot** | | | | | | |
| Favored (%) | 96.84 | 96.770 | 96.480 | 96.870 | 95.76 | 97.010 |
| Allowed (%) | 3.15 | 3.220 | 3.490 | 3.130 | 4.21 | 2.970 |
| Disallowed (%) | 0.01 | 0.010 | 0.020 | 0.000 | 0.02 | 0.020 |
| C-beta deviations (0.25 Å) | 0.000 | 0.000 | 0.000 | 0.000 | 0.000 | 0.000 |
| CaBLAM outliers (%) | 1.820 | 1.870 | 1.900 | 1.770 | 1.95 | 1.780 |
| EMRinger Score | 2.47 | 2.68 | 2.70 | 2.78 | 2.56 | 2.47 |

**Supplemental Table S2, continued. 3D reconstruction and model refinement statistics (2/3)**

| **Structure** | closed state 6 | closed state 7 | closed state 8 | closed state 9 | closed state 10 | closed state 11 |
|---|---|---|---|---|---|---|
| EM Databank Accession ID | EMD-40489 | EMD-40486 | EMD-40485 | EMD-40487 | EMD-40484 | EMD-40490 |
| Protein Data Bank Accession ID | 8SHN | 8SHF | 8SHE | 8SHG | 8SHD | 8SHO |
| **Symmetry imposed** | | | | | | |
| 3D classification | C1 | C1 | C1 | C1 | C1 | C1 |
| 3D refinement | C1 | C1 | C1 | C1 | C1 | C1 |
| Final particles | 100,823 | 37,284 | 112,259 | 120,668 | 84,821 | 34,066 |
| **Map resolution (Å)** | | | | | | |
| FSC 0.143 (unmasked) | 3.4 | 3.7 | 3.4 | 3.4 | 3.5 | 3.7 |
| FSC 0.143 (masked, corrected) | 2.8 | 3.0 | 2.8 | 2.8 | 2.9 | 3.0 |
| **Model Refinement** | | | | | | |
| Initial model used for CCT (PDB code) | 7NVM | | | | | |
| Initial model used for G$\beta_5$ (AlphaFold protein structure database) | AF-O14775-F1 | | | | | |
| Initial model used for PhLP1 (AlphaFold protein structure database) | AF-Q13371-F1 | | | | | |
| Map correlation coefficient | 0.890 | 0.870 | 0.890 | 0.880 | 0.880 | 0.880 |
| **Model composition** | | | | | | |
| Non-hydrogen atoms | 65,907 | 67,779 | 66,232 | 67,666 | 66,069 | 67,765 |
| Protein residues | 8,566 | 8,803 | 8,609 | 8,787 | 8,588 | 8,796 |
| Ligands (ADP, $AlF_x$, $Mg^{2+}$) | 16 | 16 | 16 | 16 | 16 | 16 |
| **R.m.s. deviations** | | | | | | |
| Bond lengths (Å) | 0.004 | 0.002 | 0.002 | 0.003 | 0.002 | 0.005 |
| Bond angles (˚) | 0.571 | 0.500 | 0.506 | 0.526 | 0.504 | 0.623 |
| **Validation** | | | | | | |
| MolProbity score | 1.560 | 1.590 | 1.540 | 1.610 | 1.510 | 1.660 |
| Clashscore | 6.620 | 7.240 | 6.560 | 6.990 | 6.700 | 7.360 |
| Poor rotamers (%) | 0.000 | 0.000 | 0.000 | 0.000 | 0.000 | 0.000 |
| **Ramachandran plot** | | | | | | |
| Favored (%) | 96.840 | 96.820 | 96.980 | 96.550 | 97.250 | 96.200 |
| Allowed (%) | 3.160 | 3.160 | 3.010 | 3.420 | 2.730 | 3.750 |
| Disallowed (%) | 0.000 | 0.020 | 0.010 | 0.020 | 0.020 | 0.050 |
| C-beta deviations (0.25 Å) | 0.000 | 0.000 | 0.000 | 0.000 | 0.000 | 0.000 |
| CaBLAM outliers (%) | 1.770 | 1.910 | 1.780 | 1.790 | 1.710 | 1.900 |
| EMRinger Score | 2.96 | 2.41 | 2.83 | 2.84 | 3.07 | 2.75 |

**Supplemental Table S2, continued. 3D reconstruction and model refinement statistics (3/3)**

| **Structure** | closed state 12 | closed state 13 | closed state 14 | closed state 15 | best PhLP1 class | opened state |
|---|---|---|---|---|---|---|
| EM Databank Accession ID | EMD-40491 | EMD-40492 | EMD-40494 | EMD-40461 | EMD-40481 | EMD-40439 |
| Protein Data Bank Accession ID | 8SHP | 8SHQ | 8SHT | 8SGL | 8SH9 | 8SFE |
| **Symmetry imposed** | | | | | | |
| 3D classification | C1 | C1 | C1 | C1 | C1 | C1 |
| 3D refinement | C1 | C1 | C1 | C1 | C1 | C1 |
| Final particles | 43,246 | 107,887 | 33,752 | 69,572 | 175,869 | 104,907 |
| **Map resolution (Å)** | | | | | | |
| FSC 0.143 (unmasked) | 3.7 | 3.4 | 3.8 | 3.5 | 3.2 | 4.1 |
| FSC 0.143 (masked, corrected) | 3.0 | 2.9 | 3.0 | 2.9 | 2.7 | 3.4 |
| **Model Refinement** | | | | | | |
| Initial model used for CCT (PDB code) | 7NVM | | | | | 6QB8 |
| Initial model used for G$\beta_5$ (AlphaFold protein structure database) | AF-O14775-F1 | | | | | N/A |
| Initial model used for PhLP1 (AlphaFold protein structure database) | AF-Q13371-F1 | | | | | N/A |
| Map correlation coefficient | 0.84 | 0.89 | 0.83 | 0.890 | 0.83 | 0.8 |
| **Model composition** | | | | | | |
| Non-hydrogen atoms | 68,052 | 68,037 | 68,382 | 67,432 | 67,324 | 60,512 |
| Protein residues | 8,834 | 8,831 | 8,873 | 8,762 | 8,742 | 7,881 |
| Ligands (ADP, $AlF_x$, $Mg^{2+}$) | 16 | 16 | 16 | 16 | 16 | 16 |
| **R.m.s. deviations** | | | | | | |
| Bond lengths (Å) | 0.003 | 0.005 | 0.002 | 0.004 | 0.003 | 0.003 |
| Bond angles (˚) | 0.522 | 0.565 | 0.517 | 0.551 | 0.524 | 0.651 |
| **Validation** | | | | | | |
| MolProbity score | 1.60 | 1.63 | 1.61 | 1.590 | 1.57 | 2.120 |
| Clashscore | 6.76 | 6.59 | 7.31 | 6.560 | 6.23 | 15.360 |
| Poor rotamers (%) | 0.000 | 0.000 | 0.00 | 0.000 | 0.000 | 0.000 |
| **Ramachandran plot** | | | | | | |
| Favored (%) | 96.49 | 96.12 | 96.73 | 96.530 | 96.50 | 93.390 |
| Allowed (%) | 3.50 | 3.84 | 3.25 | 3.470 | 3.5 | 6.55 |
| Disallowed (%) | 0.01 | 0.03 | 0.02 | 0.000 | 0.000 | 0.060 |
| C-beta deviations (0.25 Å) | 0.000 | 0.000 | 0.000 | 0.000 | 0.000 | 0.000 |
| CaBLAM outliers (%) | 1.88 | 1.95 | 1.86 | 1.820 | 1.88 | 3.210 |
| EMRinger Score | 2.74 | 2.9 | 2.74 | 2.74 | 3.40 | 0.85 |
